# ATGL-mediated lipid droplet lipolysis promotes collective migration in Drosophila

**DOI:** 10.64898/2026.01.06.697938

**Authors:** Israel J. Wipf, Emma J. Lowden, Julia R. Dorale, Kennedy F. Godfredsen, Michelle S. Giedt, Tina L. Tootle

## Abstract

While lipid droplets (LDs), dynamic organelles central to lipid and energy homeostasis, are implicated in cancer cell migration, their roles during collective cell migration remain unknown. We use *Drosophila* border cell migration as an *in vivo* model of invasive, collective cell migration to dissect the roles of LDs and the conserved LD lipase, Adipose Triglyceride Lipase (ATGL). Border cell LDs undergo dynamic changes and decrease in volume by the end of migration. Loss of ATGL increases LD volume, whereas border cell overexpression depletes LDs. Loss, border cell knockdown or overexpression of ATGL delays migration and blocks delamination. Further, loss of ATGL disrupts border cell mitochondria – it alters morphology, reduces membrane potential and increases reactive oxygen species. These results demonstrate that tight regulation of lipid mobilization from LDs, including for energy production, drives delamination and collective migration. Our findings not only have the potential to inform how cancer cells exploit LDs to promote their invasive behaviors but also highlight the crucial role of LDs in migration during development, hinting at their broader significance in diverse migratory contexts.

## Introduction

Collective cell migration – the movement of groups of cells to precise locations – is essential for embryonic development and wound healing but is co-opted during cancer metastasis (Carmona-Fontaine *et al*., 2008; Cheung *et al*., 2013; Riahi *et al*., 2015; Padmanaban *et al*., 2019). Emerging evidence supports that most metastases are seeded by collectively, rather than individually, migrating cancer cells, as the presence of circulating tumor cell clusters in the bloodstream is associated with worse prognosis (Murlidhar *et al*., 2017; Wang *et al*., 2017a). Therefore, there is a critical need to understand the mechanisms governing collective cell migration to facilitate wound healing and combat diseases such as cancer.

In addition to coordination between cells, collective cell migration requires the successful coordination of many intracellular processes. As a group, collectively migrating cells must dynamically alter their adhesions, remodel their membranes, and rearrange their cytoskeletal networks (Cheung *et al*., 2013; Mayor and Etienne-Manneville, 2016; De Pascalis and Etienne-Manneville, 2017). Migration is therefore an energy-intensive process, and an important question is how migrating cells, including metastatic cancer cells, produce sufficient ATP to fuel their invasion and migration (Zanotelli *et al*., 2018; Mosier *et al*., 2021). While proliferative cancer cells rely preferentially on glycolysis for ATP production (Warburg *et al*., 1927), accumulating evidence supports that metastasizing cancer cells are dependent on mitochondrial lipid oxidation to fuel their invasive and migratory behavior (LeBleu *et al*., 2014; Cunniff *et al*., 2016; Lin *et al*., 2019). Consequently, the synthesis, storage, and utilization of lipids are key processes that must be properly coordinated for efficient cell movement.

Lipid storage and utilization across species is regulated by lipid droplets (LDs) (Walther and Farese, 2012; Welte, 2015; Olzmann and Carvalho, 2019). LDs are dynamic organelles that are comprised of a phospholipid monolayer surrounding stored neutral lipids, including triglycerides and sterol esters. Thus, LDs act as safe reservoirs for toxic fatty acids (FAs) which could otherwise cause membrane damage and disrupt normal cellular functions (Lee *et al*., 1994; Nguyen *et al*., 2017). LDs are formed at the endoplasmic reticulum (ER) (Kalantari *et al*., 2010; Jacquier *et al*., 2011). ER-resident enzymes synthesize neutral lipids, which are then deposited between the leaflets of the ER bilayer. Beyond a threshold concentration, the neutral lipids demix, coalesce into an oil lens and expand to bud off from the ER as a nascent LD (Hamilton *et al*., 1983; Thiam and Foret, 2016; Hegaard *et al*., 2022). As nascent LDs mature, they form a reservoir that both protects the cell from FAs and supplies other organelles with lipids for diverse metabolic needs.

As metabolic needs arise, FAs can be rapidly mobilized from LDs by lipolysis, the hydrolysis of triglyceride into glycerol and free FAs (Walther and Farese, 2012; Welte, 2015; Olzmann and Carvalho, 2019). Canonical LD lipolysis is mediated by a cascade of neutral lipases, including adipose triglyceride lipase (ATGL), hormone-sensitive lipase (HSL), and monoglyceride lipase (MGL) (Zimmermann *et al*., 2004). Once mobilized from LDs, free FAs can serve as building blocks for membrane lipids and signaling molecules or as oxidative substrates for β-oxidation and adenosine triphosphate (ATP) production (Walther and Farese, 2012; Welte, 2015; Olzmann and Carvalho, 2019).

Dysregulated LD homeostasis is a shared feature of several types of cancer (Krahmer *et al*., 2013; Cruz *et al*., 2020; Zhang *et al*., 2022). Increasing evidence supports a role for LD dysregulation in promoting the invasive and migratory behaviors that mediate cancer metastasis (Nieman *et al*., 2011; Balaban *et al*., 2017; Wang *et al*., 2017b; Rozeveld *et al*., 2020; Andolino *et al*., 2025). For example, LD accumulation in KRAS-driven pancreatic adenocarcinoma cells primes the cells for later invasion and migration, where the excess FAs can be broken down by mitochondrial β-oxidation to generate ATP (Rozeveld *et al*., 2020). While these and other data support a role for LDs in cancer cell invasion and migration, the majority of these studies were performed using cultured cells *in vitro*. Comparatively little is known about the role of LDs during *in vivo* cell migrations, including whether LDs have a conserved role in promoting collective migration outside of the context of cancer.

To address these knowledge gaps, we use the migration of border cells during *Drosophila* oogenesis as an *in vivo* model of invasive, collective cell migration. Each *Drosophila* ovary consists of 15-20 strands of developing follicles or egg chambers called ovarioles. There are 14 stages of follicle development. Within each follicle there are 16 germline-derived cells – 15 nurse cells which support the developing oocyte – surrounded by a population of ∼650 somatic follicle cells (Giedt and Tootle, 2023).

During Stage 9 (S9), 6-8 follicle cells are specified as the border cells, detach from the anterior follicular epithelium, and collectively migrate posteriorly between the nurse cells to the oocyte. As a model system, border cell migration recapitulates many aspects of other collective cell migrations. Like both epithelial cancer cells and neural crest cells, the border cells are induced from stationary cells that undergo dramatic changes to gain motility, including changes in polarity, adhesions, and cytoskeletal organization (Montell, 2003; Montell *et al*., 2012; Saadin and Starz-Gaiano, 2016).

Once induced, the border cells must detach from an epithelium and delaminate as a cluster. Epithelial cancers (e.g., breast, colon, and prostate cancer) and melanomas also detach from an epithelium and disseminate as clusters (Hegerfeldt *et al*., 2002; Cui and Yamada, 2013; Cheung *et al*., 2016; Zajac *et al*., 2018). Finally, the border cells invade an adjacent tissue – a hallmark of metastatic cancer cells – as the somatically-derived border cells invade and migrate through the adjacent germline-derived nurse cells toward the oocyte. Many mutants have been identified that delay or prevent border cell migration, often revealing fundamental mechanisms of collective cell migration that are conserved in other systems (Montell, 2003; Montell *et al*., 2012; Saadin and Starz-Gaiano, 2016).

In this study, we sought to determine the role of LDs during border cell migration. Our analysis of migration under normal physiological conditions reveals that LDs are present within the border cell cluster and undergoing dynamic changes in both number and volume as invasion and migration proceeds. Specifically, we observe smaller LDs in late migration, suggesting LD lipids are being mobilized during migration. This finding led us to explore the role of the lipase ATGL (*Drosophila* Brummer, Bmm) in border cell migration. Our data supports the model that ATGL promotes lipid mobilization from LDs; ATGL loss increases LD volume within the border cell cluster, whereas border cell specific overexpression of ATGL depletes LDs within the border cells. Importantly, genetic loss, border cell knockdown or overexpression of ATGL leads to migration defects, including delayed migration and a failure to detach (i.e. delaminate) from the anterior follicular epithelium. These findings suggest that the border cells are sensitive to the amount of ATGL present and that successful delamination and invasive, collective migration depends on proper storage and utilization of LD-stored neutral lipids. Our data supports the model that these lipids (free FAs) are trafficked to the mitochondria for β-oxidation; loss of ATGL alters mitochondrial morphology and distribution, decreases membrane potential relative to organelle mass, and accumulates ROS in the mitochondria in the border cell cluster.

To our knowledge, this work is the first to demonstrate that tight regulation of mobilization and utilization of stored lipids from LDs is essential for successful *in vivo* delamination and migration. Given the ubiquity of LDs as lipid storage organelles, precise regulation over LD storage, mobilization, and utilization is likely a conserved requirement in diverse migratory contexts across species.

## Results

### Lipid droplets are present and dynamic within the border cells during migration

During *Drosophila* mid-oogenesis, LDs accumulate within the nurse cells and are subsequently transferred to the oocyte (Buszczak *et al*., 2002; Teixeira *et al*., 2003; Parra-Peralbo and Culi, 2011). However, whether LDs are present in the border cells was not known. To address this question, we labeled both the border cell cluster and LDs. We detect LDs specifically within the border cell cluster before and throughout migration (Fig. 1A-E). In general, pre-migratory border cells appear to contain fewer, large LDs, while BCs later in migration contain numerous small LDs throughout the cluster (Fig. 1B-E). To assess these apparent differences in the LDs, we quantified the number and size of LDs specifically within the border cell cluster before and throughout migration using Imaris software (see Materials and Methods for details).

**FIGURE 1.**
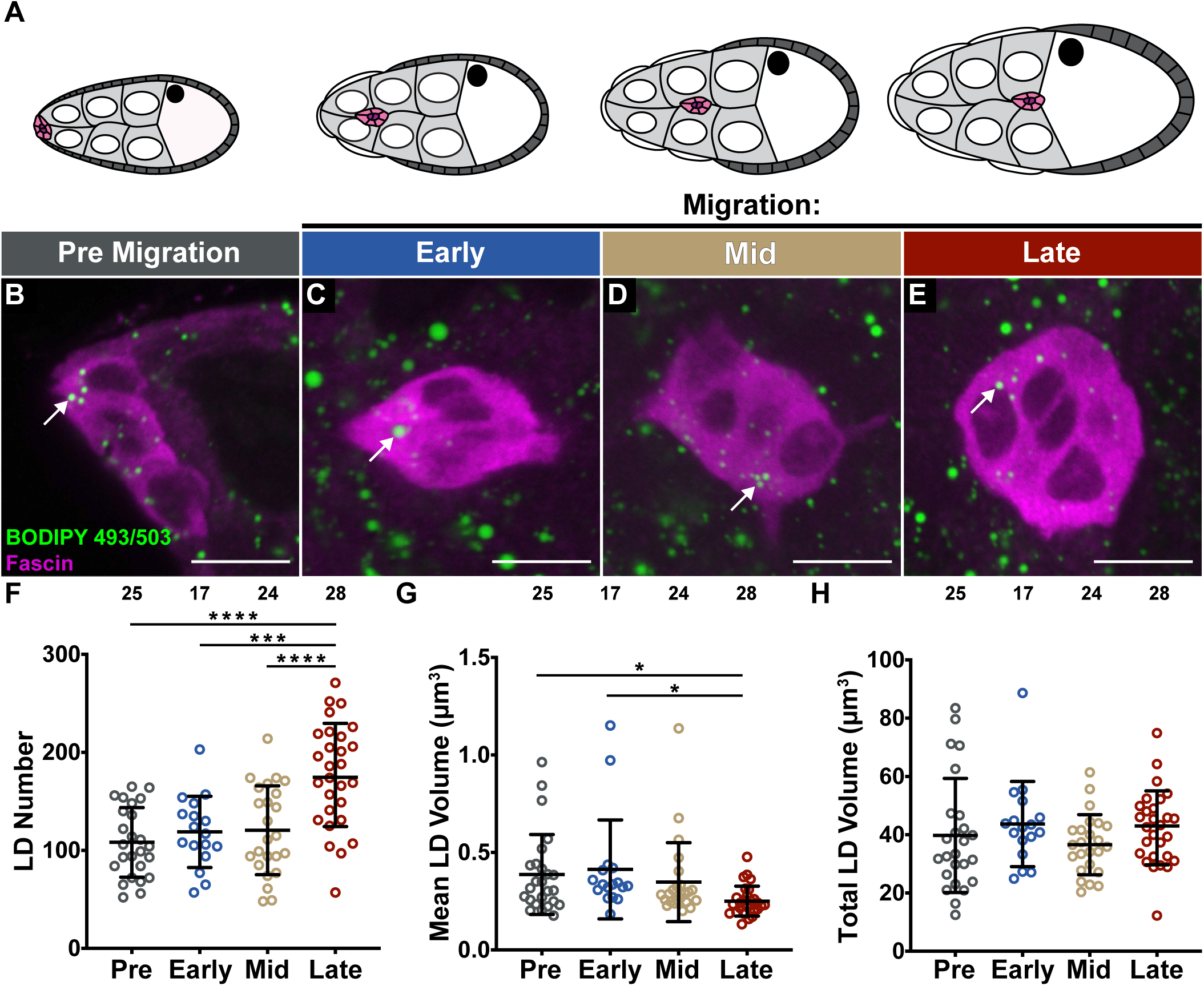
Lipid droplets (LDs) are present and dynamic in the border cells throughout migration. **(A).** Schematic of border cell migration during S9 of oogenesis. The border cells (pink), polar cells (purple), outer follicle cells (dark grey), stretch follicle cells (white), nurse cells (light grey with white nuclei), and oocyte (white with black nucleus) are diagramed. **(B-E).** Maximum projections of 3 confocal slices of wild-type (*yw*) border cells at the indicated stage of migration **(A)** stained for LDs (BODIPY 493/503) in green and Fascin in magenta. White arrows indicate examples of LDs in the border cell clusters. Images brightened by 30% to increase clarity. Black boxes placed behind panel labels to increase legibility. Scale bars = 10 μm. **(F-H).** Quantification of LD number **(F)**, mean LD volume **(G)**, and total LD volume **(H)** in the border cell cluster at the indicated stage of migration. Circle = individual border cell cluster, n = number of border cell clusters, line = mean, error bars = SD, and ns > 0.05, * p < 0.05, *** p < 0.001, **** p < 0.0001, one-way analysis of variance (ANOVA) with Tukey multiple comparison test. During S9, the border cells delaminate from the anterior follicular epithelium and migrate as a collective between the nurse cells to the nurse cell-oocyte boundary; at the same time the follicle grows in size **(A)**. LDs are present within the border cell cluster prior to and throughout their migration to the oocyte **(B-E)**. LD number within the border cell cluster increases during late migration relative to earlier stages of migration **(F)**. Mean LD volume decreases in the border cell cluster during late migration relative to pre and early migration **(G)**, while total LD volume in the border cell cluster remains unchanged throughout migration **(H)**.

We find the number of LDs (∼115) remains consistent within the border cell cluster until late migration, whereupon LD number (∼177) increases significantly compared to pre-migration (Fig. 1F; *p* < 0.0001), early migration (*p* < 0.001), and mid-migration (*p* < 0.0001). This result could indicate that later in migration there is an increase in LD synthesis, a decrease in LD utilization, or both. Supporting that stored lipids being released from within LDs throughout migration contribute to these changes, LD volume decreases at late migration (Fig. 1G; ∼0.25 µm³) relative to both pre migration (∼0.39 µm³; *p* < 0.05) and early migration (∼0.41 µm³; *p* < 0.05). However, there is no detectable change in total lipid volume in the border cells across migration (Fig. 1H;∼40 µm³); this result is consistent with the observations that border cell LDs are increasing in number yet decreasing in size throughout migration. To rule out the possibility that these changes in LD number and volume are due to changes in the border cell cluster size during migration, we measured the size of the border cell cluster at each point in migration and found there are no significant differences (Fig. S1). Together, these data reveal that LDs are not only present within the border cell cluster before and throughout migration but are undergoing dynamic changes in number and size that may contribute to the processes of detachment, invasion, and migration.

### ATGL regulates lipid droplet volume in the border cell cluster

As LDs are present and dynamic in the border cells, we next investigated whether LD lipolysis is important for border cell migration. A key lipolysis enzyme is ATGL (Gronke *et al*., 2005). To determine if ATGL regulates LDs in the border cell cluster, we analyzed LDs in *ATGL* null mutants. Loss of ATGL results in visibly larger LDs within the border cell cluster (Movie 1), which can be observed at multiple points throughout migration (Fig. 2A-H). This result suggests that lipolysis and lipid utilization from LDs within the border cells during migration are decreased when ATGL is lost.

**FIGURE 2.**
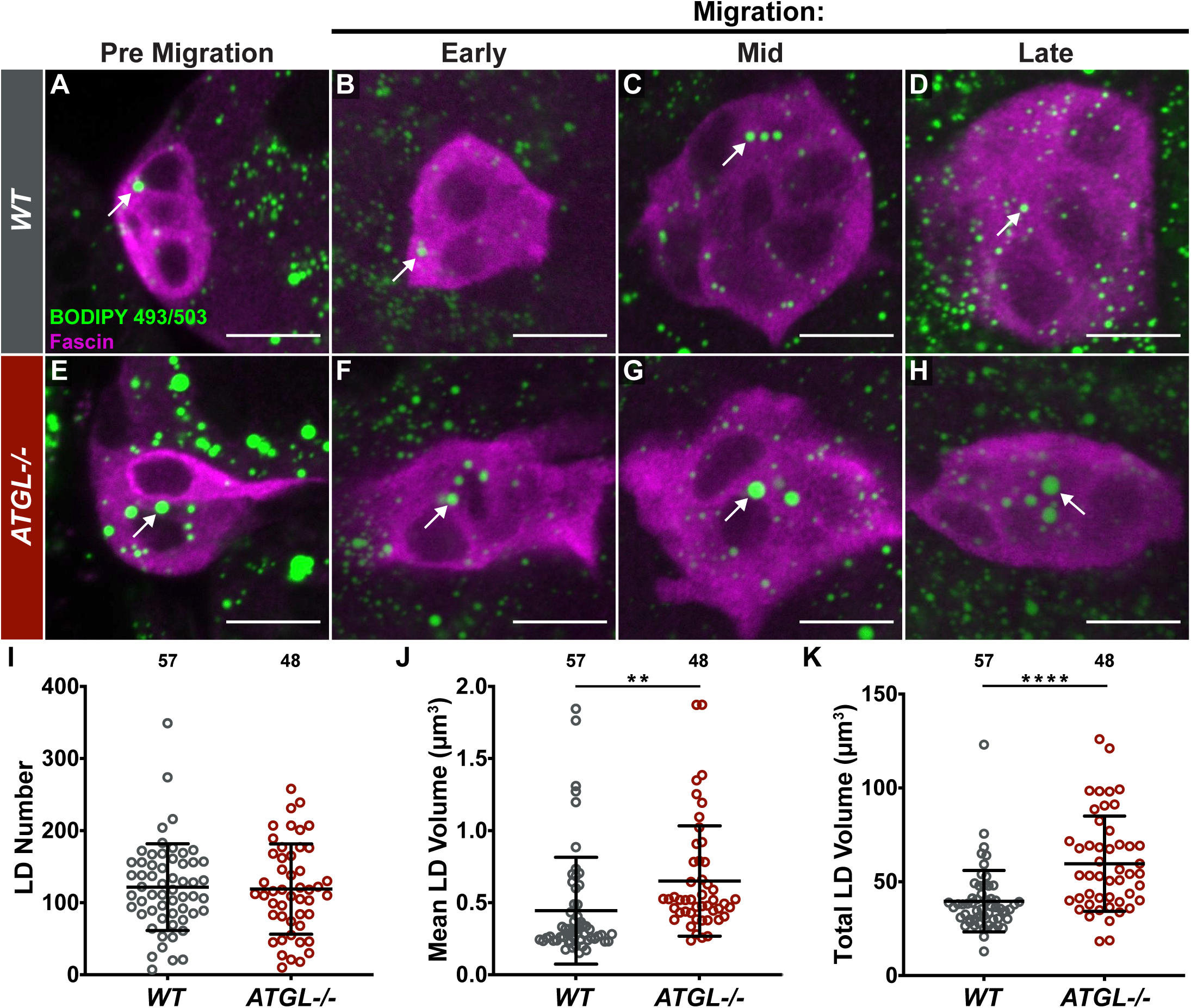
ATGL regulates lipid droplet (LD) volume in the border cell cluster**. (A-H).** Maximum projections of 3 confocal slices of S9 follicles with the border cells at the indicated stage of migration stained for LDs (BODIPY 493/503) in green and Fascin (border cells) in magenta. White arrows indicate examples of LDs in the border cell cluster. Images brightened by 30% to increase clarity (panels **B** and **H** brightened 50%). Black boxes placed behind channel labels to increase legibility. Scale bars = 10 μm. **(A-D)** *wild-type* (*yw*). **(E-H)** *ATGL-/-* (*bmm^1^/bmm^1^*). **(I-K).** Quantification of LD number **(I)**, mean LD volume **(J)**, and total LD volume **(K)** in the border cell clusters for the indicated genotypes. Circle = individual border cell cluster, n = number of border cell clusters, line = mean, error bars = SD, and ns > 0.05, ** p < 0.01, **** p < 0.0001, unpaired t-test, two-tailed. Loss of ATGL results in larger LDs within the border cell cluster relative to wild-type throughout migration **(A-H)**. Loss of ATGL does not affect LD number in the border cell cluster **(I)** but increases both mean LD volume **(J)** and total LD volume **(K)** within the border cell cluster.

Quantification of the LDs reveals that while the number of LDs within the border cell cluster remains unchanged (Fig. 2I), loss of ATGL increases mean LD volume (Fig. 2J; *p* < 0.01). Loss of ATGL also increases total lipid volume (Fig. 2K; *p* < 0.0001) in the border cell cluster. This result may be due, in part, to an increased border cell cluster size in *ATGL* mutants (Fig. S2A). Alternatively, excess lipid storage may cause the expanded border cell cluster size. These data suggest ATGL is a critical regulator of LD lipolysis in the border cell cluster.

### ATGL is required for border cell detachment and on-time migration

We next asked whether ATGL loss affects border cell invasion and migration. Using *ATGL* mutants, we first assessed border cell cluster morphology. Despite the increased size of the border cell cluster, we find loss of ATGL does not alter the length of the cluster (Fig. S2B). Next, to assess the timing of border cell migration during S9, we compare the border cell cluster’s position relative to the position of the outer follicle cells (Fig. 3A). In wild-type follicles, the border cell cluster remains in-line with the outer follicle cells throughout migration (Fig. 3B-B’). We find loss of ATGL delays border cell migration, as the cluster remains anterior to the outer follicle cells (Fig. 3C-C’). To quantify border cell migration, we measure the distance the border cell cluster has migrated away from the anterior end of the follicle (termed border cell distance) and divide it by the distance of the outer follicle cells from the anterior end of the follicle (termed follicle cell distance). This calculation generates a metric we term the migration index (Fig. 3A; (Fox *et al*., 2020; Lamb *et al*., 2020; Mellentine *et al*., 2023)). A migration index of ∼1 indicates on-time migration, while values less than 1 or values greater than 1 indicate delayed or accelerated migration, respectively. Using this method, the migration index in *ATGL* mutants (MI = 0.637; *p* < 0.0001) is significantly lower than wild-type controls (Fig. 3D; MI = 0.911). These data indicate ATGL is required for on-time border cell migration.

**FIGURE 3.**
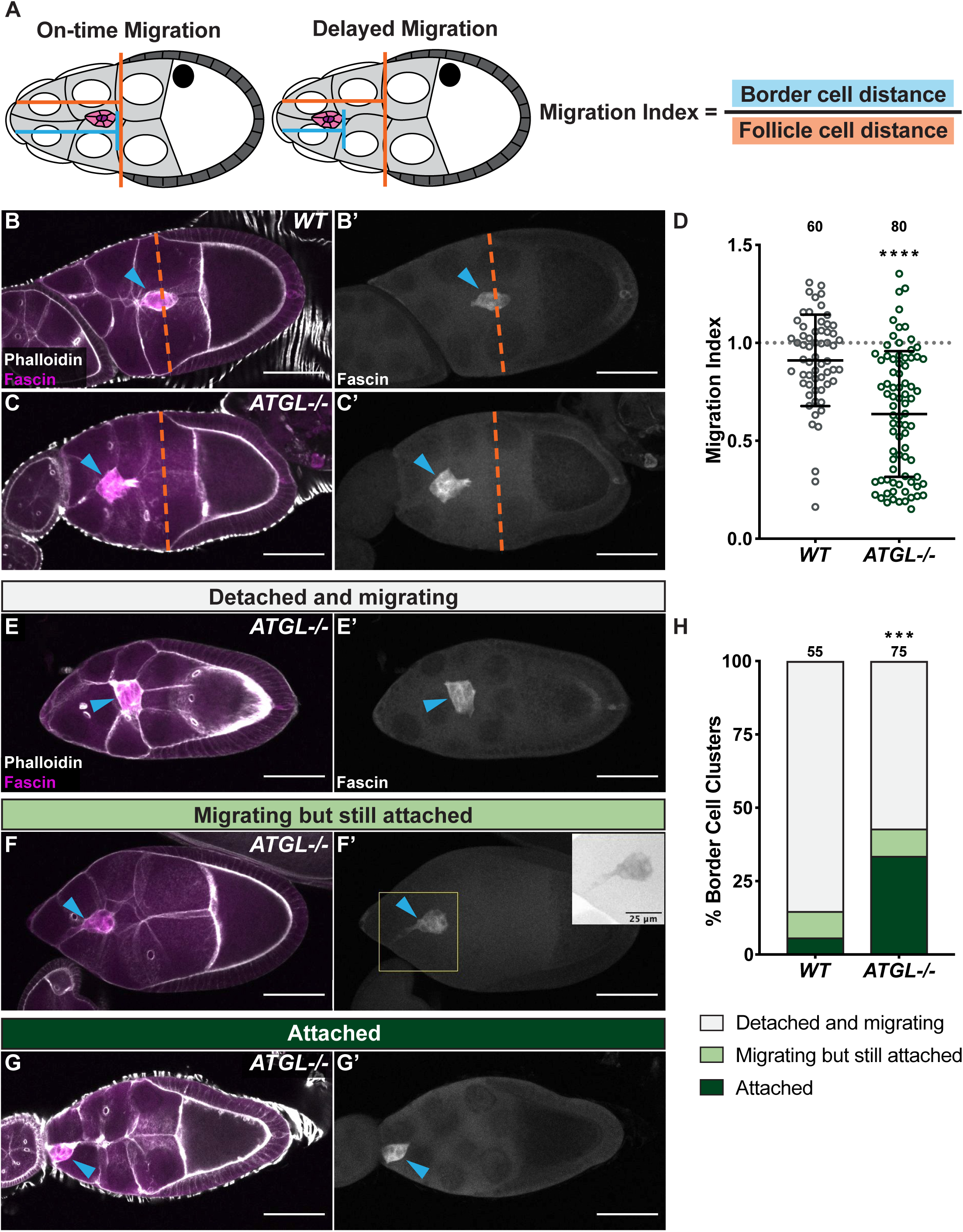
ATGL is required for on-time border cell migration and detachment. **(A).** Schematic of on-time (left) and delayed (right) border cell migration, and the formula for calculating the Migration Index. In each schematic, anterior is on the left and posterior is on the right. The border cells (pink), polar cells (purple), outer follicle cells (dark grey), stretch follicle cells (white), nurse cells (light grey with white nuclei), and oocyte (white with black nucleus) are diagramed. The Migration Index is calculated by dividing the distance the border cells have migrated (blue line) by the distance of the outer follicle cells from the anterior end of the follicle (orange line). **(B, C).** Maximum projections of 3 confocal slices of S9 follicles stained for Fascin (border cells) in magenta and F-actin (Phalloidin) in white. Blue arrowheads indicate the border cell cluster, and orange dashed lines indicate the anterior edge of the outer follicle cells. Images brightened by 30% to increase clarity. Black boxes placed behind panel and channel labels to increase legibility. Scale bars = 50 μm. **(B, B’)** *wild-type* (*yw*). **(C, C’)** *ATGL-/-* (*bmm^1^/bmm^1^*). **(D).** Graph of migration index for the indicated genotypes. Circle = individual border cell cluster, n = number of border cell clusters. The dotted line = on-time border cell migration, solid black lines = means, error bars = SD, and ns > 0.05, **** p < 0.0001, unpaired *t*-test, two-tailed. **(E-G).** Maximum projections of 3 confocal slices of *ATGL-/-* S9 follicles showing border cell clusters that are detached from the follicular epithelium and migrating **(E)**, migrating but still attached **(F)**, and attached **(G)**. White arrowheads indicate the border cell cluster. **(H).** Quantification of border cell detachment defects for the indicated genotypes. Border cell clusters categorized as detached and migrating (light grey), migrating but still attached (light green), or attached (dark green). n = number of border cell clusters, *** p < 0.001, Pearson’s Chi-squared test. The timing of border cell migration can be assessed during S9 by comparing the distance the border cells have migrated relative to the position of the outer follicle cells **(A)**. In wild-type follicles, the migrating border cell cluster must detach from the follicular epithelium and migrate; during migration the cluster remains in-line with the outer follicle cells **(A, left; D, H)**. Loss of ATGL delays border cell migration during S9 **(C, D)** and impairs detachment of the border cell cluster from the follicular epithelium **(E-H)**.

While quantifying the migration index, we observed many *ATGL* mutant follicles with extremely low migration index values. This observation led us to investigate the role of ATGL in the process of border cell detachment or delamination. The border cell cluster is specified from nonmotile, epithelial follicle cells. Thus, before invasion and migration can proceed, the border cell cluster must first detach from the anterior follicular epithelium. In contrast to wild-type follicles, we find that in *ATGL* mutants the border cell clusters often fail to detach from the anterior follicular epithelium during S9. To quantify defects in border cell detachment, we classified mid-to late-S9 follicles into three categories: those with border cell clusters that are fully detached and migrating (Fig. 3E-E’), migrating but have a tail that remains attached to the anterior follicular epithelium (Fig. 3F-F’), or attached to the anterior follicular epithelium (Fig. 3G-G’). Loss of ATGL resulted in 33.3% of border cell clusters remaining attached to the anterior follicular epithelium during S9, compared to just 5.5% in wild-type controls (Fig. 3H; *p* < 0.001). Correspondingly, only 57.3% of ATGL mutant clusters were detached and migrating, compared to 85.5% in wild-type controls (Fig. 3H; *p* < 0.001). Loss of ATGL did not impact the number of clusters that were migrating but still attached (9% vs. 9.3% in wild-type controls). These data indicate ATGL is required for border cell delamination, suggesting that the *ATGL* S9 migration delay may be due, at least partially, to an initial failure to detach from the anterior follicular epithelium.

### ATGL RNAi knockdown in the border cells exhibits only minor changes in LDs

Given the increased LD volume within the border cell clusters of *ATGL* null mutants, we asked whether we could detect changes in border cell cluster LDs when depleting ATGL using a border cell-specific RNAi approach. To this end, we used the UAS/GAL4 system to knockdown ATGL by RNAi in the border cells and analyzed border cell LDs. In contrast to the *ATGL* mutant phenotype (Fig. 2), RNAi knockdown of ATGL in the border cells does not alter border cell LD volume (Fig. 4A-C, E-F). However, knockdown of ATGL in the border cells increases LD number (Fig. 4D) compared to the GAL4 only control (p = 0.0528) but not compared to the UAS only control. The difference in border cell LD phenotypes between *ATGL* mutants and border cell-specific ATGL RNAi could reflect differences in RNAi knockdown efficiency, ATGL protein stability, the timing of GAL4-driven induction, or some combination of these factors. It is also possible that the border cell LD phenotype observed in *ATGL* mutants is due to loss of ATGL in either the outer follicle cells – which give rise to the border cells – and/or in the nurse cells, the substrate of the border cell cluster’s migration. Further experiments are required to distinguish between these possibilities.

**FIGURE 4.**
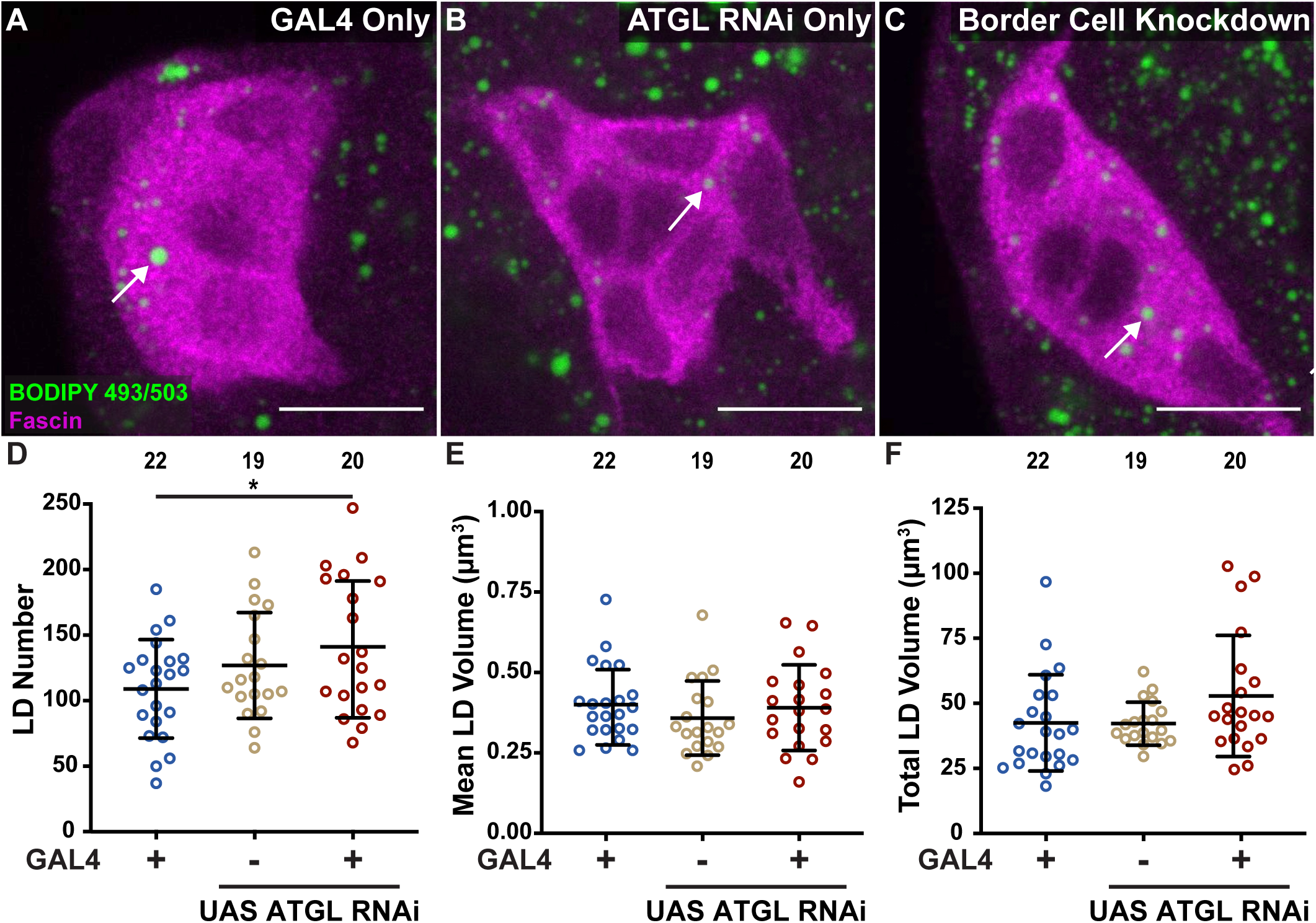
Knockdown of ATGL in the border cells increases LD number but does not alter LD volume. **(A-C).** Maximum projections of 3 confocal slices of S9 follicles stained for LDs (BODIPY 493/503) in green and Fascin (border cells) in magenta. White arrows indicate examples of LDs in the border cell cluster. Images brightened by 50% to increase clarity. Black boxes placed behind panel and channel labels to increase legibility. Scale bars = 10 μm. **(A)** GAL4 control (*slbo GAL4/+*). **(B)** ATGL RNAi control (*UAS bmm RNAi/+*). **(C)** Border cell knockdown of ATGL (*slbo GAL4/UAS bmm RNAi*). **(D-F).** Quantification of LD number **(D)**, mean LD volume **(E)**, and total lipid volume **(F)** in the border cell clusters for the indicated genotypes. Circle = individual border cell cluster, n = number of border cell clusters, line = mean, error bars = SD, * p<0.05, one-way analysis of variance (ANOVA) with Tukey multiple comparison test. Border cell knockdown of ATGL does not alter LD volume **(A-C, E, F)** but increases LD number in the border cells **(D)** compared to the GAL4 only control.

### ATGL is required in the border cells for detachment and on-time migration

We next sought to determine where ATGL is required to promote border cell detachment and on-time migration. Given that ATGL regulates LD lipolysis in a cell-autonomous manner, and our finding that ATGL regulates LDs within the border cell cluster (Fig. 2), we hypothesized ATGL is required in the border cells. To test this, we again used the UAS/GAL4 system to knockdown ATGL by RNAi in the border cells. As expected, knockdown of ATGL in the border cells delays border cell migration (Fig. 5A-D; MI = 0.575) compared to the GAL4 only (MI = 0.929; *p* < 0.0001) and RNAi only (MI = 0.834; *p* < 0.001) controls. Furthermore, like the mutant phenotype, RNAi knockdown of ATGL in the border cells results in significant detachment defects. Border cell knockdown of ATGL resulted in 39% of border cell clusters remaining attached to the anterior follicular epithelium during S9 (Fig. 5E), compared to just 12.5% and 10% in the GAL4 only (*p* < 0.05) and RNAi only (*p* < 0.01) controls, respectively. Additionally, knockdown of ATGL in the border cells increases the number of border cells that are migrating but still attached (12.5%) compared to the GAL4 only (2.5%; *p* < 0.05) and RNAi only (2.5%; *p* < 0.01) controls. Only 55.5% of border cell clusters were detached and migrating under the ATGL knockdown condition, whereas the majority of the GAL4 only (85%; *p* < 0.05) and RNAi only (87.5%; *p* < 0.01) control border cell clusters were detached and migrating. Together, these data reveal ATGL is required specifically in the border cells for border cell delamination and on-time migration.

**FIGURE 5.**
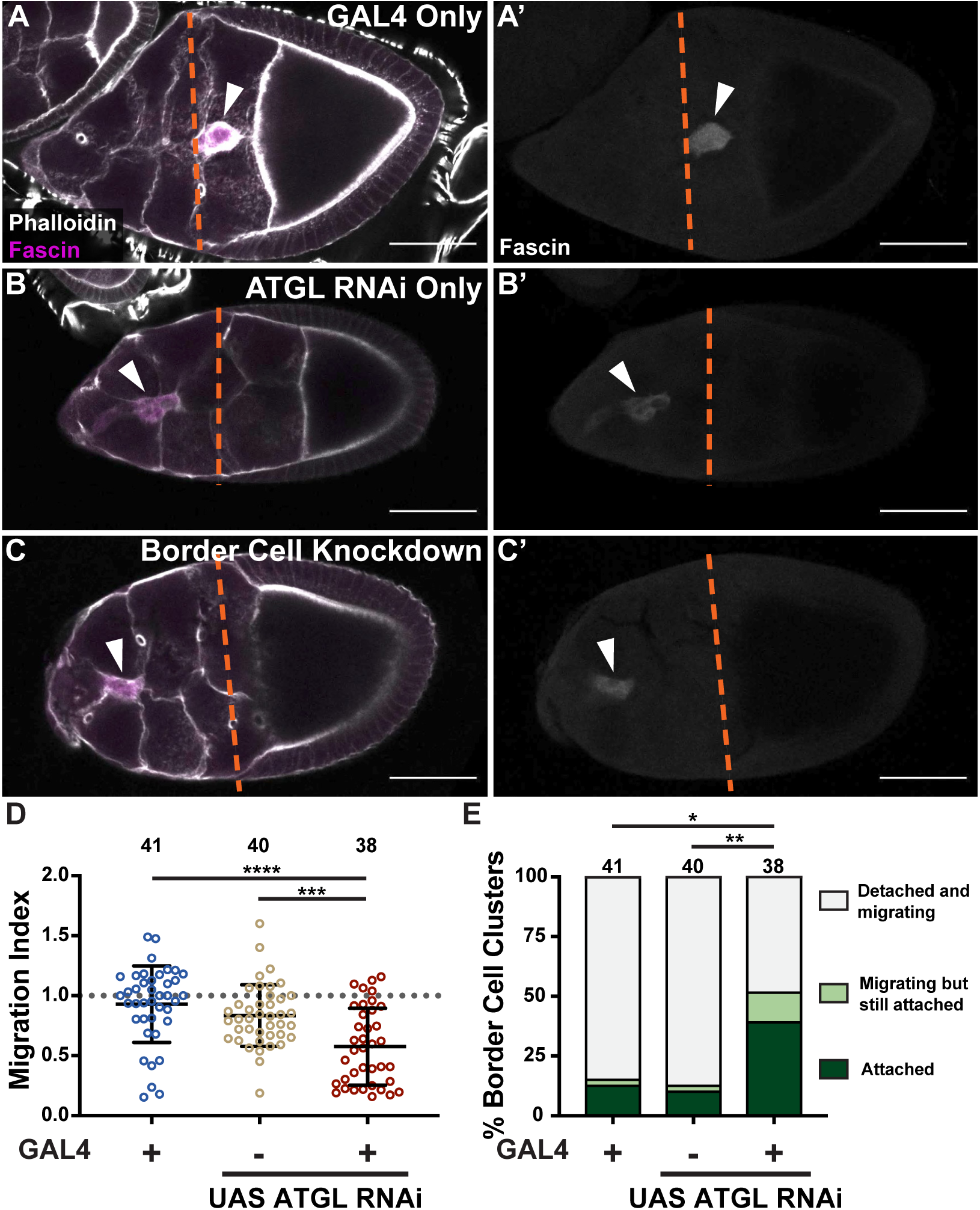
ATGL is required in the border cells for detachment and on-time migration. **(A-C).** Maximum projections of 3 confocal slices of S9 follicles stained for Fascin (border cells) in magenta and F-actin (Phalloidin) in white. White arrowheads indicate the border cell cluster, and orange dashed lines indicate the anterior edge of the outer follicle cells. Images brightened by 50% to increase clarity. Black boxes placed behind panel and channel labels to increase legibility. Scale bars = 50 μm. **(A, A’)** GAL4 control (*slbo GAL4/+*). **(B, B’)** ATGL RNAi control (*UAS bmm RNAi/+*). **(C, C’)** Border cell knockdown of ATGL (*slbo GAL4/UAS bmm RNAi*). **(D).** Graph of migration index for the indicated genotypes. Circle = individual border cell cluster, n = number of border cell clusters. The dotted line = on-time border cell migration, solid black lines = means, error bars = SD, and ns > 0.05, *** p < 0.001, **** p < 0.0001, one-way analysis of variance (ANOVA) with Tukey multiple comparison test. **(E).** Quantification of border cell detachment defects for the indicated genotypes. Border cell clusters categorized as detached and migrating (light grey), migrating but still attached (light green), or attached (dark green). *n* = number of border cell clusters, and ns > 0.05, * p < 0.05, ** p < 0.01, Pearson’s Chi-squared test with Monte Carlo simulation. Border cell knockdown of ATGL delays border cell migration during S9 **(A-D)** and impairs detachment of the border cell cluster from the follicular epithelium **(E)**.

### ATGL overexpression in the border cells depletes lipid droplets

A previous study revealed that overexpression of the lipase HSL depletes LD stores and decreases invasive potential in multiple pancreatic cancer cell lines (Rozeveld *et al*., 2020). Given our findings that ATGL regulates border cell LDs (Fig. 2) and is required in the border cells for detachment and on-time migration (Figs. 3 and 5), we asked whether the border cells might be similarly sensitive to the amount of ATGL present. To test this, we used the UAS/GAL4 system to overexpress ATGL in the border cells. While border cell LDs of GAL4 only controls appear normal (Fig. 6A), UAS ATGL only control border cells appear moderately depleted of LDs (Fig. 6B).

**FIGURE 6.**
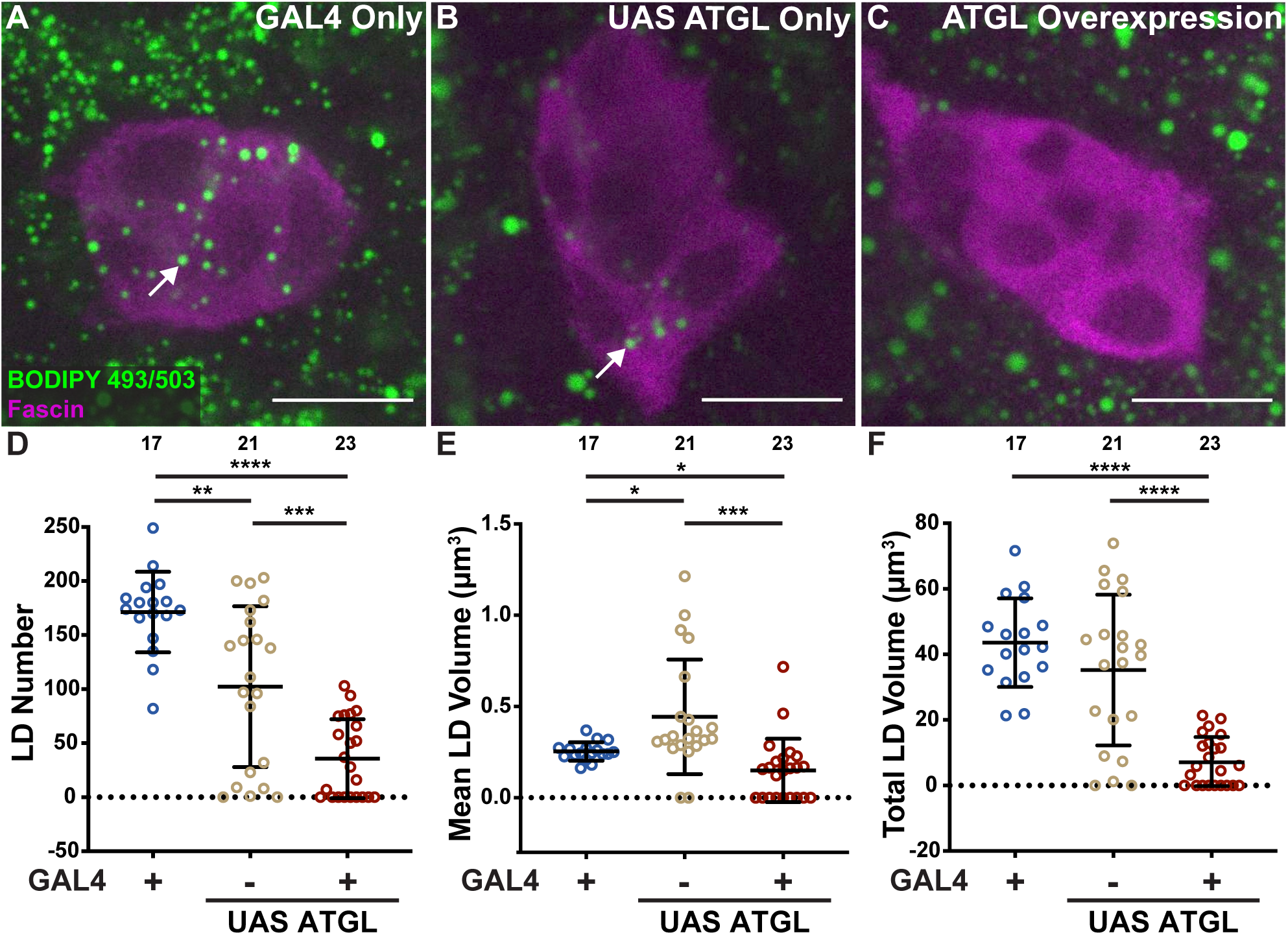
Overexpression of ATGL in the border cells depletes LDs. **(A-C).** Maximum projections of 3 confocal slices of S9 follicles stained for LDs (BODIPY 493/503) in green and Fascin (border cells) in magenta. White arrows indicate examples of LDs in the border cell cluster. Images brightened by 60% to increase clarity. Black boxes placed behind channel labels to increase legibility. Scale bars = 10 μm. **(A)** GAL4 control (*slbo GAL4/+*). **(B)** UAS ATGL control (*UAS bmm/+*). **(C)** Border cell overexpression of ATGL (*slbo GAL4/UAS bmm*). **(D-F).** Quantification of LD number **(D)**, mean LD volume **(E)**, and total lipid volume **(F)** in the border cell clusters for the indicated genotypes. Circle = individual border cell cluster, n = number of border cell clusters, line = mean, error bars = SD, and ns > 0.05, * p < 0.05, ** p < 0.01, *** p < 0.001, **** p < 0.0001, one-way analysis of variance (ANOVA) with Tukey multiple comparison test. Overexpression of ATGL in the border cells decreases LD number **(A-D)**, moderately decreases mean LD volume **(E)**, and strikingly depletes total LD volume **(F)** in the border cells.

Strikingly, however, border cell overexpression of ATGL strongly depletes border cell LDs (Fig. 6C). Quantification reveals, as the images suggest, fewer LDs in UAS ATGL only control border cell clusters compared to the GAL4 only control (∼102 compared to ∼171, respectively; p < 0.001). One possible explanation for this result is that the UAS-ATGL construct may be expressing the UAS transgene in the absence of GAL4. To test this possibility, we analyzed border cell cluster LDs in either a wild-type or *ATGL* heterozygous background combined with UAS ATGL (no GAL4). We reasoned that if the reduction of LDs observed in UAS ATGL control clusters is in fact due to increased ATGL protein, then genetically reducing ATGL may rescue the phenotype. Indeed, we find that genetic reduction of ATGL restores the number of border cell cluster LDs back to normal levels (Fig. S3). Regardless of the leaky UAS ATGL expression in the absence of GAL4, we find that border cell overexpression of ATGL dramatically decreases LD number in the border cells (Fig. 6D; ∼36) compared to both the GAL4 only (∼171; *p* < 0.0001) and UAS ATGL only (∼102; *p* < 0.001) controls. Further, border cell overexpression of ATGL moderately decreases mean LD volume in the border cells (Fig. 6E; ∼0.16 µm³) compared to the GAL4 only (∼0.25 µm³; *p* < 0.05) and UAS ATGL only (∼0.43 µm³; *p* < 0.001) controls, and dramatically decreases total lipid volume in the border cells (Fig. 6F; ∼7.29 µm³) compared to the GAL4 only (∼43.59 µm³; *p* < 0.0001) and UAS ATGL only (∼ 35.23 µm³; *p* < 0.0001) controls. This near loss of border cell cluster LDs when ATGL is overexpressed could be due to a decrease in LD biogenesis, an increase in mobilization of lipids from LD, or both.

Given ATGL’s known lipase function, however, it is likely that ATGL overexpression is excessively releasing FAs from triglyceride stores within LDs, leading to LD shrinkage and eventual depletion. Together, these data support that ATGL levels and activity must be tightly regulated in the border cells to control LD number and size.

### ATGL overexpression in the border cells delays detachment and migration

Given the striking changes in LDs when ATGL is overexpressed in the border cells, we next assessed how ATGL overexpression in the border cells affects border cell delamination and migration. As before, we used the UAS/GAL4 system to overexpress ATGL in the border cells. We find ATGL overexpression in the border cells delays border cell migration (Fig. 7A-D; MI = 0.696) compared to the GAL4 only (MI = 1.118; *p* < 0.0001) and UAS only (MI = 0.917; *p* < 0.05) controls. Additionally, we observed detachment defects similar to those observed under loss or reduction of ATGL. Border cell ATGL overexpression resulted in 37% of border cell clusters remaining attached to the anterior follicular epithelium during S9 (Fig. 7E), compared to only 6% and 17% in the GAL4 only (*p* < 0.001) and UAS only (*p* < 0.05) controls, respectively. Because the severity of the detachment and migration defects correlates with the extent of LD depletion observed within each genotype (Fig. 6), these findings support the model that proper neutral lipid storage and utilization are essential for both delamination and on-time migration, and loss of neutral lipid stores within the border cell cluster impairs these processes.

**FIGURE 7.**
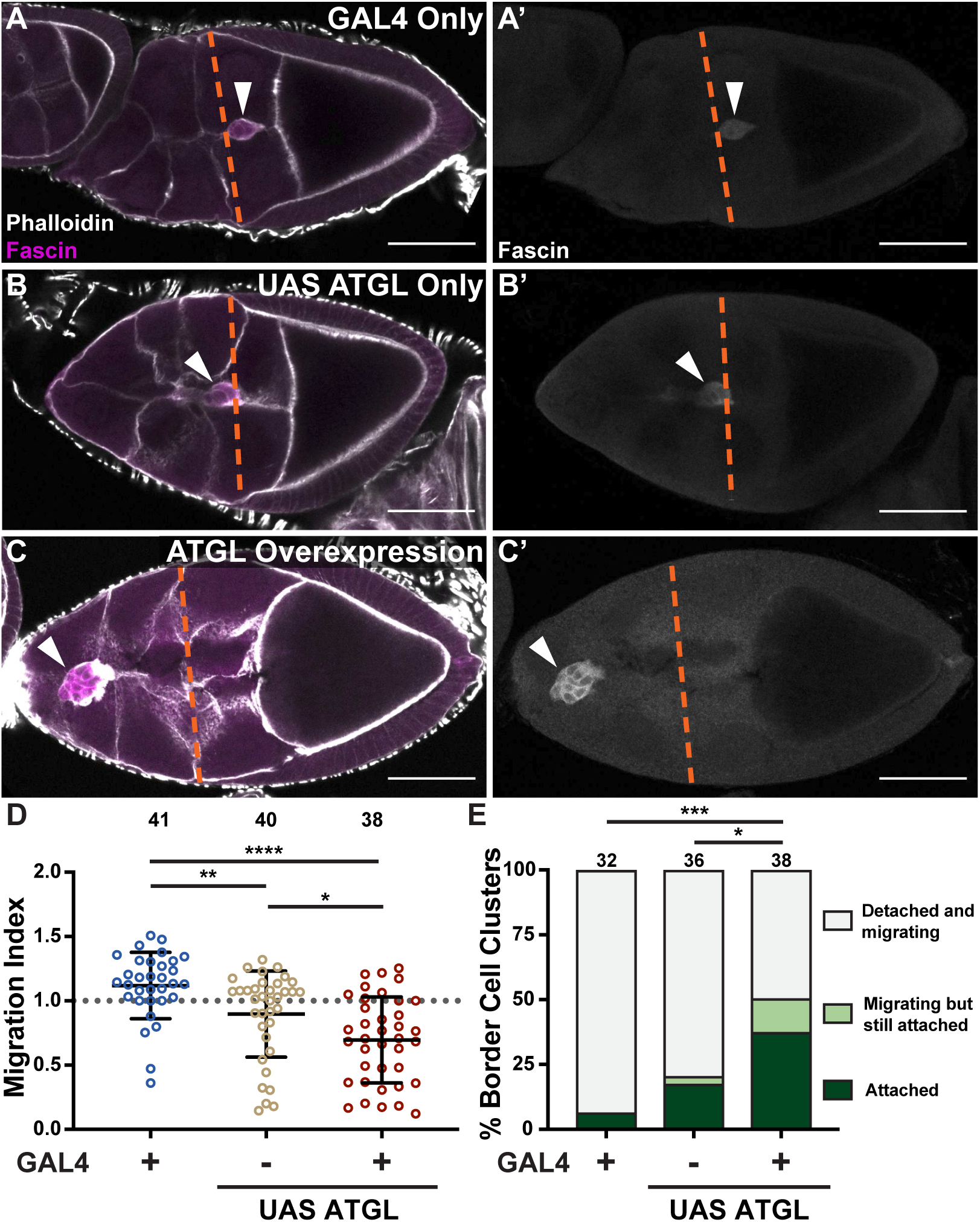
Overexpression of ATGL in the border cells impairs detachment and delays migration. **(A-C).** Maximum projections of 3 confocal slices of S9 follicles stained for Fascin (border cells) in magenta and F-actin (Phalloidin) in white. White arrowheads indicate the border cell cluster, and orange dashed lines indicate the anterior edge of the outer follicle cells. Images brightened by 50% to increase clarity. Black boxes placed behind panel labels to increase legibility. Scale bars = 50 μm. **(A, A’)** GAL4 control (*slbo GAL4/+*). **(B, B’)** UAS ATGL control (*UAS bmm/+*). **(C, C’)** Border cell overexpression of ATGL (*slbo GAL4/UAS bmm*). **(D).** Graph of migration index for the indicated genotypes. Circle = individual border cell cluster, n = number of border cell clusters. The dotted line = on-time border cell migration, solid black lines = means, error bars = SD, and * p < 0.05, ** p < 0.01, **** p < 0.0001, one-way analysis of variance (ANOVA) with Tukey multiple comparison test. **(E).** Quantification of border cell detachment defects for the indicated genotypes. Border cell clusters categorized as detached and migrating (light grey), migrating but still attached (light green), or attached (dark green). *n* = number of border cell clusters, and ns > 0.05, * p < 0.05, *** p < 0.001, Pearson’s Chi-squared test with Monte Carlo simulation. Border cell overexpression of ATGL delays border cell migration during S9 **(A-D)** and impairs detachment of the border cell cluster from the follicular epithelium **(E)**.

### Loss of ATGL disrupts mitochondrial morphology and function in the border cells

Following ATGL-mediated lipolysis, how do mobilized FAs promote border cell delamination and on-time migration? There are many possible mechanisms, as mobilized FAs could serve as precursors for membrane lipids and/or signaling molecules, or as substrates for energy production.

First, we tested the possibility that ATGL-mediated lipolysis is releasing the FA arachidonic acid for prostaglandin (PG) production as it does during S10B of *Drosophila* oogenesis (Giedt *et al*., 2023). Previously, our lab has shown that PG signaling is required for on-time border cell migration, and that loss of PG synthesis delays migration and disrupts cluster morphology (Fox *et al*., 2020; Mellentine *et al*., 2023). To test whether ATGL’s control over border cell migration is dependent on PG signaling, we performed a dominant genetic interaction assay between ATGL and dCOX1, the *Drosophila* cyclooxygenase-like enzyme responsible for all PG synthesis (Tootle and Spradling, 2008). We can perform this assay because heterozygosity for the individual mutations (ATGL−/+ or dCOX1−/+) does not appreciably effect border cell migration (Fig. S4). If ATGL acts upstream of PG synthesis, then the double heterozygotes should exhibit delayed migration. We find that co-reduction of both ATGL and dCOX1 does not delay border cell migration, as the border cells from double heterozygotes of *ATGL* and *dCOX1* migrate on time (Fig. S4). This result suggests that FAs mobilized by ATGL are acting through a mechanism other than PG production to promote border cell migration. Consistent with this, loss of PG signaling does not impact border cell detachment (Fox *et al*., 2020; Mellentine *et al*., 2023).

Given that cell migration is an energy-intensive process, we next sought to address the possibility that mobilized FAs are provided to the mitochondria for β-oxidation and energy production. Supporting this possibility, a previous study found that proper regulation of mitochondrial fission and fusion dynamics is essential for border cell migration (Qu *et al*., 2022). Additionally, LDs are often physically and functionally linked to mitochondria, and perturbations in one organelle can have detrimental effects on the other (Enkler and Spang, 2024; Fan and Tan, 2024).

Therefore, we next sought to determine if loss of ATGL affects border cell mitochondria. We labeled mitochondria in *ATGL* null mutants and found mitochondrial morphology is altered in the border cells. Compared to wild-type mitochondria which appear to be evenly dispersed throughout the cytoplasm, mitochondria of *ATGL* null border cells appear to cluster in large aggregates (Fig. S5). This mitochondrial clustering is indicative of mitochondrial dysfunction and may reflect impaired energy production in the border cell cluster.

Given the changes in mitochondrial morphology observed in *ATGL* null border cells, we next asked whether loss of ATGL impairs mitochondrial function. A fundamental indicator of mitochondrial health is the maintenance of an inner membrane potential, which is essential for ATP production through oxidative phosphorylation. To assess mitochondrial membrane potential, we employed MitoView Fix 640, a mitochondrial stain whose localization depends on mitochondrial polarization. We find that MitoView Fix 640 colocalizes to mitochondria in the border cells (Fig. 8A-A”), and that loss of ATGL does not significantly change the intensity of the staining (Fig. 8B-C). To use this staining to assess mitochondrial function, we normalized mitochondrial membrane potential to total mitochondrial mass by calculating the ratio of polarization-dependent mitochondrial signal (MitoView Fix 640) to polarization-independent signal (ATP-β). We find loss of ATGL significantly increases the maximum intensity of ATP-β staining (Fig. 8D; *p* < 0.001) and decreases the ratio of polarization-dependent to polarization-independent staining (Fig. 8E; *p* < 0.01). This result suggests that *ATGL* mutant border cells have reduced overall mitochondrial membrane potential, consistent with reduced energy production.

**FIGURE 8.**
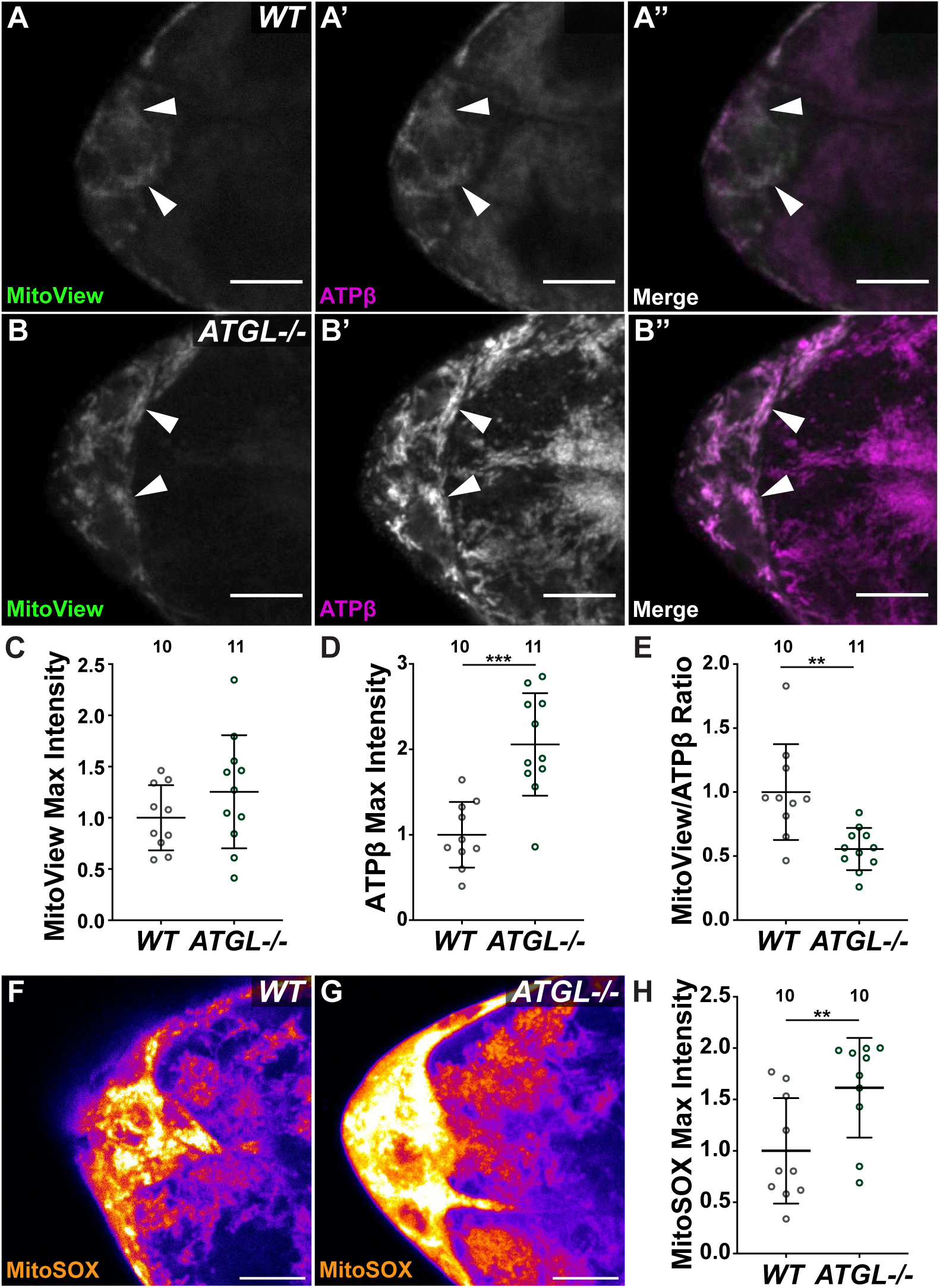
Loss of ATGL disrupts mitochondrial function in the border cells. **(A-B).** Maximum projections of 3 confocal slices of S9 follicles stained for total mitochondria (ATPβ) and oxidatively active mitochondria (MitoView). White arrowheads indicate instances of colocalization between ATPβ and MitoView in the border cell cluster. Images brightened by 50% to increase clarity. Black boxes placed behind panel and channel labels to increase legibility. Scale bars = 10 μm. **(A-A’’)** *wild-type* (*yw*). **(B-B’’)** *ATGL-/-* (*bmm^1^/bmm^1^*). **(C-E).** Quantification of maximum fluorescence intensity of MitoView **(C)**, maximum fluorescence intensity of ATPβ **(D)**, and ratio of MitoView to ATPβ signal **(E)** in the border cell cluster for the indicated genotypes. Circle = individual border cell cluster, n = number of border cell clusters, line = mean, error bars = SD, and ns > 0.05, ** p < 0.01, *** p < 0.001, unpaired t-test, two-tailed. **(F, G).** Maximum projections of 3 confocal slices of S9 follicles stained for MitoSOX. A Fire LUT was applied using ImageJ to show the levels of MitoSOX fluorescence. Panels F and G were not brightened. Black boxes placed behind panel labels to increase legibility. Scale bars = 10 μm. **(F)** *wild-type* (*yw*). **(G)** *ATGL-/-* (*bmm^1^/bmm^1^*). **(H).** Quantification of maximum fluorescence intensity of MitoSOX signal in the border cell cluster for the indicated genotypes. Loss of ATGL disrupts mitochondrial function in the border cells. Loss of ATGL results in reduced membrane potential relative to total mitochondria **(A-E)**, and increased ROS production in the border cells **(F-H)**.

Given that mitochondrial mass fails to provide a proportional increase in total membrane potential in *ATGL* mutants, we investigated if there is oxidative stress. We utilized the indicator MitoSOX to specifically detect mitochondrial superoxide, a byproduct of respiration that is toxic at high concentrations and a primary member of the reactive oxygen species (ROS) family. Consistent with mitochondrial dysfunction, loss of ATGL triggers significant increase in MitoSOX fluorescence in the border cell cluster (Fig. 8F-8H; p < 0.01). Together these data support a role for ATGL in maintaining border cell mitochondrial function and health, as its loss alters mitochondrial morphology, impairs membrane potential, and increases oxidative stress. All of these defects could be due to impaired FA β-oxidation.

Last, we asked whether FA import into the mitochondria is required for border cell migration. Carnitine palmitoyltransferase 1 (CPT1; *Drosophila* Withered, Whd) is a rate-limiting enzyme for FA β-oxidation that transports long-chain FAs into the mitochondria (Strub *et al*., 2008). Using *CPT1* null mutants, we assessed the role of CPT1 in border cell migration. Loss of CPT1 results in delayed border cell migration (Fig. S6A-C; MI=0.295 compared to MI= 1.002, *p* < 0.0001) and strikingly, results in 89.3% of border cell clusters remaining attached to the anterior follicular epithelium during S9 (Fig. S6D; *p* < 0.0001), compared to just 6.6% in the wild-type. This phenotype is similar, but more severe, than what is observed when ATGL is lost (Fig. 3). Together, these findings strongly suggests that FA import into the mitochondria, including FAs released from LD stores by ATGL, is essential for border cell delamination and therefore, migration.

## Discussion

Using *Drosophila* border cell migration as a model, we provide the first evidence that the regulated mobilization and utilization of stored lipids from LDs is essential for successful *in vivo* delamination and collective migration during development (Fig. 9). We find that the border cells store neutral lipids in LDs prior to and during their migration, and that these LDs undergo changes in size and number as migration proceeds. Further, we identify the neutral lipase ATGL as an important regulator of LD lipolysis in the border cells, as loss of ATGL significantly increases the size of LDs within the border cell cluster. Through both mutant analysis and cell-specific RNAi, we demonstrate that ATGL is required specifically in the border cells for both on-time migration and for the initial detachment (delamination) of the border cell cluster from the anterior follicular epithelium. While our data supports that ATGL-mediated lipolysis promotes border cell delamination and migration, it must be carefully regulated. Excess release of stored lipids could lead to lipotoxicity or other cellular dysfunctions, especially if those lipids are later unavailable when needed. Indeed, overexpression of ATGL in the border cells almost entirely depletes LD stores within the border cell cluster, and results in both failed delamination and delayed migration. Evidence supports that tight regulation of ATGL-mediated LD lipolysis is critical for regulating mitochondria function, as loss of ATGL results in altered mitochondrial morphology and distribution, decreased membrane potential relative to organelle mass, and dramatic ROS accumulation in the border cell cluster. We speculate that these mitochondrial defects are due to decreased FA β-oxidation as loss of CPT1, the enzyme that transports FAs into the mitochondria for β-oxidation (Houten *et al*., 2016), phenocopies loss of ATGL – failed delamination and delayed migration. Together these data lead to the model that ATGL-dependent LD lipolysis is tightly regulated during border cell migration to, at least in part, ensure that FAs are available for β-oxidation in the mitochondria to provide energy to fuel delamination and migration.

**FIGURE 9.**
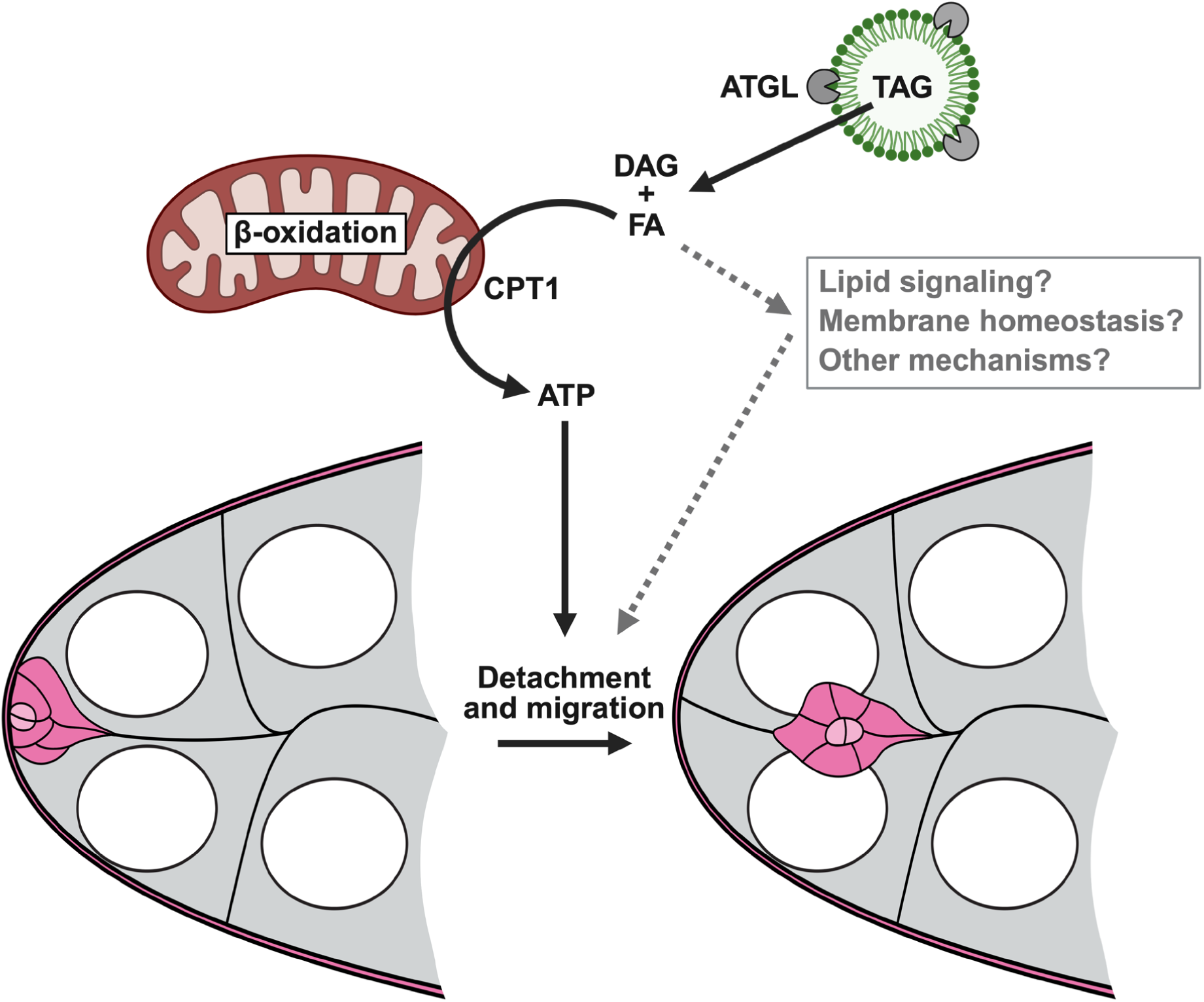
ATGL-mediated lipolysis promotes border cell delamination and migration by supporting mitochondrial function. Schematic (created with BioRender.com) summarizing our findings. LDs (green) are present in the border cell cluster (pink) and are dynamic during migration. The lipase ATGL (grey notched circles) cleaves LD-stored triglycerides (TAG) to generate free FAs and diacylglycerol (DAG). Our data supports one function of these FAs is to be transported by CPT1 into the mitochondria (brown) for β-oxidation and the production of ATP. This ATP likely fuels the delamination and migration of the border cells. Our data do not exclude the possibility that the ATGL-released FAs have other functions within the border cells, including lipid signaling and membrane homeostasis.

### Lipolysis facilitates border cell migration

Our data reveals that mobilization and utilization of stored lipids from LDs is critical for the delamination and collective migration of the border cells. LDs are present in the border cells throughout migration but are larger and fewer in number pre-and during migration than at the end of migration (Fig. 1). LD size and number are regulated by LD biogenesis, fusion, and utilization. Large LDs are indicative of older LDs or a cellular state promoting LD storage, while smaller LDs indicate LD biogenesis or high lipid utilization, resulting in LD shrinkage (Walther and Farese, 2012; Welte, 2015; Olzmann and Carvalho, 2019). We speculate that biogenesis and utilization both occur during border cell migration to maintain LD stores while fueling migration. In contrast, lipid needs shift at the end of migration, resulting in more numerous and smaller LDs – a change suggesting biogenesis and/or depletion of LD stores. Indeed, the transcription factor that drives the expression of lipid biosynthetic genes, Sterol Regulator Element-Binding Protein, is active in border cells (Sieber and Spradling, 2015).

Supporting that lipids from LDs are being utilized during border cell migration, loss of the rate limiting enzyme in triglyceride lipolysis, ATGL, increases LD size but not number (Fig. 2). The lack of an increase in LD number suggests that either small, newly formed LDs rapidly fuse with existing LDs or that there is a feedback mechanism preventing LD biogenesis when lipids are not being mobilized from LDs. Further, loss or border cell knockdown of ATGL results in both impaired delamination and delayed migration (Figs. 3 and 5), revealing this key lipolytic enzyme is critical for the initial invasion and the collective migration of the border cells. Notably, RNAi knockdown of ATGL does not result in LD changes (Fig. 4); we speculate this is due to ATGL levels being reduced but not absent due to either the timing of or extent of knockdown. It is also possible that ATGL activity within other cells also contributes to the LD phenotype observed in the border cells in the *ATGL* mutant. Finally, overexpression of ATGL in the border cells results in striking LD depletion (Fig. 6) and impaired delamination and delayed migration (Fig. 7). The similarity in the border cell migration defects between too little and too much ATGL strongly supports that ATGL levels, and likely activity, must be tightly regulated to ensure lipid utilization from LDs occurs at the right time and right level to mediate the delamination and collective migration of the border cells.

Border cell delamination is regulated by numerous factors. At the beginning of S9, the polar cells secrete the ligand Unpaired, which activates JAK/STAT signaling within the adjacent outer follicle cells (Duchek *et al*., 2001; Silver and Montell, 2001). This signaling specifies them to become border cells, turning on the expression of Slow Border Cells (Slbo), a C/EBP transcription factor, which activates downstream factors essential for delamination and migration (Montell *et al*., 1992). One factor is the cell-adhesion molecule E-Cadherin, which both facilitates adhesions within the cluster and migration by adhesions to the nurse cells (Oda *et al*., 1997; Niewiadomska *et al*., 1999). Loss of any of these components results in impaired (E-Cadherin) or complete failure to delaminate (all the others). However, a recent study uncovered distinctions in these mutants (Inaki *et al*., 2022). Specifically, loss of Jak/STAT signaling results in the border cells failing to differentiate, remaining within the follicular epithelium and being immotile, whereas when Slbo is lost the border cell cluster rounds up, the cells move back and forth, but fail to form protrusions and cannot invade (Inaki *et al*., 2022). Loss of E-Cadherin is similar to loss of Slbo, except the border cells form smaller and less stabile protrusions. We find that loss, border cell knockdown, or border cell overexpression of ATGL severely impairs border cell delamination. In these contexts, the border cells express Fascin, another downstream target of Slbo (Borghese *et al*., 2006; Wang *et al*., 2006), indicating the border cells are specified. Further, when ATGL levels are perturbed, the border cell clusters round up and some eventually migrate (Figs. 3, 5 and 7). These findings suggest that tight regulation of ATGL is not required for border cell specification but is required for their delamination.

Our data supports that the defects in delamination and subsequent migration when ATGL levels are perturbed are due, at least in part, to defective lipolysis and its downstream effects on mitochondria. Loss of ATGL results in clustered, elongated, and perhaps increased mitochondria in the border cells (Fig. S5; Fig. 8). These findings are consistent with increased mitochondrial fusion, increased mitochondrial biogenesis, or both. Mitochondrial fission is critical for border cell migration, as impairing fission by loss of Drp1 or overexpression of mitofusins delays and impairs the completion of migration (Qu *et al*., 2022). The study suggests, through *in vitro* and whole ovary analyses, that the migration defects are due to reduced mitochondrial ATP production. Supporting similar mitochondrial defects arise due to loss of ATGL, while there appears to be increased mitochondria, there is not a proportional increase in membrane potential, indicating reduced membrane potential per unit of mitochondria (Fig. 8) – a finding indicative of mitochondrial dysfunction. Consistent with this, loss of ATGL triggers mitochondrial ROS production. Supporting that this mitochondrial dysfunction is due to altered FA β-oxidation, loss of CPT1 phenocopies the delamination defects observed when ATGL levels are perturbed (Fig. S6).

Together these data support the model that ATGL-released FAs from LDs are used, at least in part, for β-oxidation within the mitochondria to produce energy necessary for the delamination and on-time migration of the border cells. Supporting this ATGL function is conserved across contexts, a recent study found that ATGL-mediated lipolysis is essential for mitochondria function, as mutating or inhibiting ATGL decreases mitochondrial-LD interaction, and disrupts mitochondria respiration and network connectivity in skeletal muscle (Gemmink et al. 2026).

One energy intensive aspect of cell invasion and migration is remodeling of the actin cytoskeleton (DeWane *et al*., 2021). Indeed, actin remodeling is key for both border cell delamination and migration (Montell, 2003; Montell *et al*., 2012). Loss of two actin binding proteins – Profilin (Ghiglione *et al*., 2018) and Fascin (Lamb *et al*., 2020) – results in delamination defects. While we don’t observe any striking changes in F-actin levels within the border cells upon altering ATGL levels, it remains likely that F-actin dynamics within the cluster are altered and contributing to the defects in delamination and migration.

### LDs, ATGL, and β-oxidation – conserved regulators of invasion and migration

The functions of ATGL, LDs and mitochondrial β-oxidation in invasive migration are likely conserved across other developmental contexts and in cancer metastasis.

Indeed, mouse migrating neural crest cells exhibit LD accumulation, although the function of these LDs has not been explored (Patel *et al*., 2015). Most of the evidence supporting roles for LDs, lipolysis, and β-oxidation in cell migration comes from *in vitro* studies on cancer cells and correlative studies on human patient samples. For example, metastatic breast cancer cells exhibit higher LD accumulation compared to non-metastatic cells; knocking down ATGL or inhibiting FA β-oxidation reduces 2D migration and 3D invasion (Wang *et al*., 2017b; Andolino *et al*., 2025). Similar findings were observed in pancreatic cancer cell lines (Rozeveld *et al*., 2020). Supporting LDs contribute to cancer progression in humans, breast cancer lung metastases from patients show LD accumulation (Andolino *et al*., 2025) and LD density positively correlates with higher Gleason scores in prostate cancer (Nardi *et al*., 2019). Thus, LDs have emerged as mediators of cancer migration, invasion, and metastasis, and are thus, biomarkers of aggressive cancer and poor patient outcomes.

The lipids mediating LD formation within the cancer cells come from both *de novo* lipogenesis in the cancer cells and uptake from the environment. Supporting that cancer cells can produce lipids *de novo*, many cancers upregulate FA synthesis enzymes, such as fatty acid synthase (FASN). For example, in metastatic breast cancer cell lines, depletion or inhibition of FASN reduces *in vitro* 3D cell invasion and survival, and *in vivo* metastasis (Andolino *et al*., 2025). FASN is overexpressed in malignant peripheral nerve sheath tumors (Patel *et al*., 2015) and high FASN levels are a cancer biomarker (Baenke *et al*., 2013; Jin *et al*., 2023). Cancer cells are often surrounded by adipocytes, and metabolic syndromes with excess lipid storage are associated with an increased risk for cancer (Lamabadusuriya *et al*., 2025). Numerous studies employing the co-culture of adipocytes and cancer cell lines – breast, ovarian, and melanoma – reveal cancer cells deplete adipocytes of lipid stores (Dirat *et al*., 2011; Nieman *et al*., 2011; Lazar *et al*., 2016; Balaban *et al*., 2017; Wang *et al*., 2017b; Clement *et al*., 2020). These lipids are then stored in LDs in the cancer cells, where they are released by lipases, including ATGL to drive CPT1-dependent mitochondrial β-oxidation to fuel proliferation, migration, and invasion (Balaban *et al*., 2017; Wang *et al*., 2017b). Thus, LDs produced from endogenous and exogenous lipids are a critical source of energy production for cancer cell migration and invasion.

Where the lipids come from for LD formation and subsequent utilization to drive border cell delamination and migration remains unknown. However, given the high concentration of LDs in the environment (the nurse cells) surrounding the border cells, it is tempting to speculate some of the lipids used to produce the border cell LDs are coming from the environment. Supporting this idea, loss of ATGL, but not border cell knockdown of ATGL, results in a striking increase in LD size. This difference could be due to a role of ATGL in releasing FAs within the nurse cells that are subsequently taken up into the outer follicle cells (prior to border cell specification) and/or the border cells during migration. Future studies exploring the roles of lipid uptake into and FA synthesis within the border cells are needed to determine the origin of the lipids within border cell LDs.

Emerging evidence suggests LD and mitochondrial localization are also critical for cell migration. For example, in pancreatic cancer cells during 2D collective migration and 3D invasion, LDs are depleted from the leading and invading cells, respectively (Rozeveld *et al*., 2020). In melanoma cells co-cultured with adipocytes, mitochondria undergo fission, and mitochondria and LDs are redistributed to cell protrusions (Clement *et al*., 2020); this mitochondrial localization is blocked by inhibiting FA β-oxidation. In ovarian cancer cells, mitochondria are enriched in lamellipodia during 3D invasion, resulting in increased ATP production to fuel actin remodeling (Cunniff *et al*., 2016). Similarly, mitochondria are trafficked to the invasive front for ATP production necessary for actin protrusions during *C. elegans* anchor cell invasion (Kelley *et al*., 2019). Finally, targeted localization of LDs regulates the migration of phytoplankton, revealing an ancient link between LD metabolism and motility (Sengupta *et al*., 2022). Together these studies reveal that not only are LD utilization and mitochondrial FA β-oxidation important for invasion and cell migration, but the localization of these organelles is critical.

As border cell migration, like all invasive, collective migrations, is highly energy dependent, a key remaining question is whether the distribution of LDs, mitochondria, and ATP production from β-oxidation is preferentially at the leading edge. Border cells regularly change which cell leads the cluster (Prasad and Montell, 2007). Thus, it is possible that energy depletion – decreased LDs and ATP production – within the leading cell drives another cell to take over as leader. Future studies, including live imaging studies of LD dynamics, are needed to address this possibility and to determine if there is polarity of LDs and mitochondria within the border cell cluster.

## Conclusion

Together our results reveal that tight regulation of ATGL-dependent release of FAs from LD stores within the border cells is critical for the delamination and on-time collective migration of the *Drosophila* border cells. We provide evidence that these FAs are used for β-oxidation, as mitochondrial morphology is altered and its function is impaired when ATGL is lost. This study is the first evidence that LDs and ATGL contribute to delamination and migration in a normal, developmental context. Given that LDs, ATGL, and β-oxidation have emerged as key drivers of cancer cell migration, invasion, and metastasis, we speculate this energy production pathway is widely used across migratory contexts.

### Study limitations

The data presented strongly supports that tight regulation of the LD lipase ATGL is necessary for on-time delamination and migration of the *Drosophila* border cells. The underlying mechanisms remain to be fully elucidated. Evidence supports that one mechanism is regulating mitochondrial function, as the mitochondria are larger and appear more prevalent, but exhibit reduced membrane potential relative to organelle mass and increased ROS when ATGL is lost; these findings are consistent with a reduction in FA β-oxidation. Future studies are needed to fully test this model. Additionally, how LDs dynamically change in number, polarity within the cluster, and in formation and utilization during migration remains unknown.

## Materials and Methods

### Reagents and resources

See Supplementary Table S1 for detailed information on the reagents used in these studies, Supplementary Table S2 for the specific genotypes used in each figure panel, and Supplementary Table S3 for all raw data reported in this study.

### Fly stocks

Fly stocks were maintained on Bloomington standard fly food (cornmeal/agar/yeast food) at 22°C except where noted. The following stocks were obtained from the Bloomington Drosophila Stock Center (BDSC): *y^1^w^1^* (BDSC, #1495), *slbo GAL4* (BDSC, #58435), *bmm^1^* (BDSC, #98123, (Gronke *et al*., 2005)), *UAS-bmm RNAi* (BDSC, #25926), *UAS-bmm* (BDSC, #76600), and *whd^1^* (BDSC, #441). Expression of the RNAi lines for follicle analyses was achieved by crossing to *slbo GAL4*, maintaining the fly crosses at room temperature (22°C), and maintaining progeny for border cell analyses at 29°C for 4-5 days. Expression of other UAS lines (*e.g., UAS-bmm*) for follicle analyses was achieved by crossing to *slbo GAL4*, maintaining the fly crosses at room temperature, and maintaining progeny for border cell analyses at 25°C for 4-5 days. Before immunofluorescence staining, newly eclosed flies were fed wet yeast paste every day for 4-5 days.

### Immunofluorescence

*Drosophila* ovaries (5-8 pairs per sample) were dissected in room temperature Grace’s insect medium (Corning). Ovaries were fixed for 10 min with 4% paraformaldehyde diluted in Grace’s medium. Samples were washed 6 times for 10 min each at room temperature in antibody wash (1X phosphate-buffered saline [PBS], 0.1% Triton X and 0.1% bovine serum albumin). The following primary antibodies were used: mouse anti-Fascin 1:50 (sn7c, Cooley, L., AB_528239; (Cant *et al*., 1994)) and mouse anti-ATP-β 1:250 (ab14730; Abcam). The anti-Fascin primary antibody was obtained from the Developmental Studies Hybridoma Bank (DSHB) which was developed under the auspices of the National Institute of Child Health and Human Development and maintained by the Department of Biology, University of Iowa (Iowa City, IA). Primary antibodies were diluted in antibody wash and incubated overnight at 4°C. The samples were washed 6 times for 10 min each at room temperature in antibody wash. Samples were then incubated with fluorescent secondary antibodies diluted 1:250 in antibody wash. Secondary antibodies used were goat anti-mouse Alexa Fluor 488 (AB_2534069; ThermoFisher Scientific) and goat anti-mouse Alexa Fluor 568 (AB_2534072; ThermoFisher Scientific). Alexa Fluor 568-or Alexa Fluor 647-conjugated Phalloidin (A12380 and A22287; Thermo Fischer Scientific) diluted 1:250 was included in both primary and secondary antibody incubations. To stain for LDs, following antibody staining, samples were incubated with BODIPY 493/503 (4,4-Difluoro-1,3,5,7,8-Pentamethyl-4-Bora-3a,4a-Diaza-s-Indacene; Invitrogen D3922) at a concentration of 1:100 in 1X PBS for 20 mins at room temperature. After LD staining, 4′,6-diamidino-2-phenylidole (DAPI; 5 mg/mL; D3571; Thermo Fischer Scientific) staining was performed at a concentration of 1:5000 in 1X PBS for 10 min at room temperature. Finally, samples were rinsed 3 times in 1X PBS and mounted on coverslips in 1 mg/mL phenylenediamine in 50% glycerol, pH 9 (Platt and Michael, 1983).

### Mitochondrial membrane potential and superoxide production staining

To assess mitochondrial membrane potential, freshly dissected ovaries were incubated in Stage 9 (S9) medium containing MitoView™ Fix 640 (200 nM, Biotium) for 90 minutes at room temperature (Prasad and Montell, 2007). S9 media consists of Schneider’s medium (Sigma-Aldrich), 0.6 x penicillin/streptomyocin (Life Technologies, SCR_008817), 0.2 mg/mL insulin (Sigma-Aldrich, SCR_008988), and 15% fetal bovine serum (Atlanta Biologicals). After MitoView staining, samples were rinsed twice in 1X PBS, then fixed and stained following the immunofluorescence protocol described above.

To assess mitochondrial superoxide production, freshly dissected ovaries were incubated in Grace’s medium containing MitoSOX™ Red (5μM, Invitrogen) for 15 minutes at room temperature. After MitoSOX staining, samples were rinsed twice in 1X PBS and fixed in 4% paraformaldehyde for 10 minutes. After fixation, samples were rinsed three times in 1X PBS and stained with DAPI (1:5000) for 10 min.

Samples were then rinsed three more times in 1X PBS and mounted on coverslips in 1 mg/mL phenylenediamine in 50% glycerol, pH 9 (Platt and Michael, 1983).

### Image acquisition and processing

Microscope images for fixed and stained *Drosophila* follicles were taken using LAS-X software (SCR_013673) on a Leica DMi8 Stellaris using a HCPLAPO CS2 20x/0.75 Dry or HCPL APO CS2 63x/1.4 Oil, or Zen software (SCR_013672) on Zeiss 700 LSM mounted on an Axio Observer.Z1 using a Plan-Apochromat 20x/0.8 M27 or EC-Plan_Neo_Fluar 40x/1.3 Oil (Carl Zeiss Microscopy). S9 follicles were identified by their size (∼150 μm–250 μm) and morphology, including the location of the outer follicle cells and the border cell cluster. The beginning of S10A was defined as when the anterior most outer follicle cells reached the nurse cell-oocyte boundary and flattened. Maximum projections, merge, rotation, cropping, and confocal image stack movie generation were performed using ImageJ software (FIJI, RRID: SCR 002285, (Abramoff *et al*., 2004)). Images were brightened as indicated in the figure legends to improve visualization. Figures were made using Illustrator (Adobe, RRID: SCR 010279).

### Quantification of border cell migration, detachment, and cluster morphology

Quantification of the Migration Index (MI) was performed as described previously (Fox *et al*., 2020; Lamb *et al*., 2020; Mellentine *et al*., 2023). Briefly, measurements of S9 follicles were performed using ImageJ software (Abramoff *et al*., 2004) on Z-stacks of follicles stained for DAPI, Fascin, and Phalloidin. A line segment was used to measure the distance in microns from the anterior end of the follicle to the leading edge of the border cell cluster; this was defined as the border cell distance. Another line segment was used to measure the distance from the anterior end of the follicle to the anterior end of the outer follicle cells: this was defined as the outer follicle cell distance. The entire follicle length was also measured along the anterior-posterior axis. The MI was calculated by dividing the border cell distance by the follicle cell distance. The length of the border cell cluster was determined by measuring the distance from the front to the rear of the border cell cluster (attached clusters were not included in cluster length analyses). The volume of the border cell cluster was quantified using the Fascin signal and the 3D surface objects tool of Imaris (version 10.2; Bitplane, Concord, MA). Data compilations and calculations were performed in Excel (Microsoft, RRID: SCR 016137), and graphs were generated and statistical analyses performed using Prism (GraphPad Software, RRID SCR 002798).

Z-stacks projection images of S9 follicles were used to analyze the position of the border cell cluster for detachment defects. S9 follicles stained for DAPI, Fascin, and phalloidin were classified into three categories: those with border cell clusters that are 1) fully detached and migrating, 2) migrating but have a tail that remains attached to the anterior follicular epithelium, or 3) attached to the anterior follicular epithelium.

Only mid-to late-S9 follicles were scored; early S9 follicles were not included in delamination quantifications. Data compilations and calculations were performed in Excel, graphs were generated using Prism, and Pearson’s Chi-squared analyses were performed using RStudio (4.5.2 (2025-10-31)).

### Lipid quantification

S9 follicles were stained and imaged on a Leica DMi8 Stellaris as described above. The resulting z-stacks were converted to Imaris files (.ims). To analyze LDs specifically in the border cell cluster, the 3D surface objects tool of Imaris (version 10.2; Bitplane, Concord, MA) was used to create a region of interest from the Fascin signal. This approach was used to create a Fascin-positive region of interest for the border cell cluster, called the border cell surface. To restrict LD signal to the border cell cluster, the BODIPY 493/503 signal was masked to the border cell surface. Following masking, the abundance and volume of LDs within the border cell cluster was quantified using the 3D surface objects tool of Imaris. LD surfaces were generated using automated thresholding (background subtraction) performed with local contrast. To resolve closely apposed objects, the split touching objects (region growing) function was enabled with a seed point diameter of 0.6 μm, followed by a morphological split.

Seed points were filtered by applying a quality threshold above an automated cutoff. The analysis settings were kept constant between genotypes. Following rendering, LD quantifications were exported as Microsoft Excel files. Each data point represents the averaged value within one border cell cluster. Data were compiled in and calculations were performed in Excel, and graphs were generated and statistical analyses performed using Prism.

### Mitochondrial quantifications

To quantify mitochondrial membrane potential, MitoView and ATP-β intensity was measured in ImageJ from single confocal slices of immunofluorescence images of fixed S9 follicles. The “straight line” function was used to draw lines between 4–7 microns at three different locations within the border cell cluster, and the highest fluorescence intensity value was measured for both MitoView and ATP-β. To calculate the MitoView to ATP-β staining ratio, the average value was calculated for the three line segments for each border cell cluster, then the MitoView value was divided by the ATP-β value, and the average was normalized to the overall wild-type average. Data were compiled in and calculations were performed in Excel, and graphs were generated and statistical analyses performed using Prism.

To quantify mitochondrial superoxide production, MitoSOX intensity was measured in ImageJ from single confocal slices of fixed S9 follicles. The “straight line” function was used to draw lines between 4–7 microns at three different locations within the border cell cluster, and the highest fluorescence intensity value was measured for MitoSOX. Lines between 4–7 microns were also used to measure the background intensity value outside of the follicle, which was subtracted. The average value was then calculated for the three line segments and the average was normalized to the overall wild-type average. Data were compiled in and calculations were performed in Excel, and graphs were generated and statistical analyses performed using Prism.

## Acknowledgements

We thank the Tootle lab for helpful discussions and careful review of the manuscript. Stocks obtained from the Bloomington Drosophila Stock Center (NIH P40OD018537) were used in this study. FlyBase (release FB2024_04) was used for information on stocks. At the University of Iowa, Information Technology Services – Research Services provided data storage support.

## Data and resource availability

All relevant data and details of resources can be found within the article and its supplementary information.

---

**Supplementary Figure 1.**
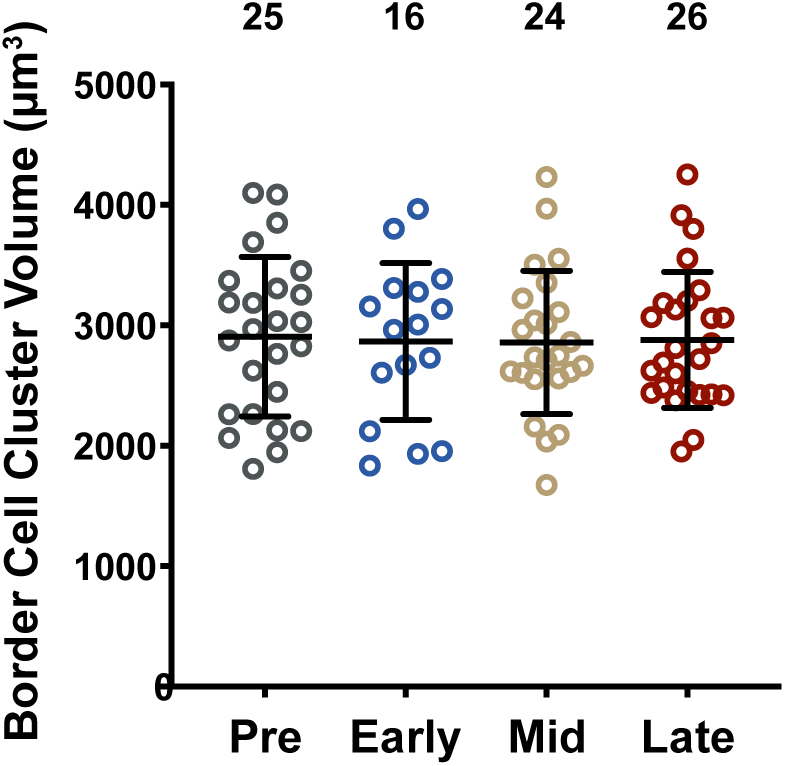
Border cell cluster volume is unchanged throughout migration. Quantification of the *wild-type* (*yw*) border cell cluster volume at the indicated stages of migration. Circle = individual border cell cluster, n = number of border cell clusters, line = mean, error bars = SD, and ns > 0.05, one-way analysis of variance (ANOVA) with Tukey multiple comparison test. In *wild-type* (*yw*) flies, the size of the border cell cluster (∼3000 μm^3^) does not change throughout migration.

**Supplementary Figure 2.**
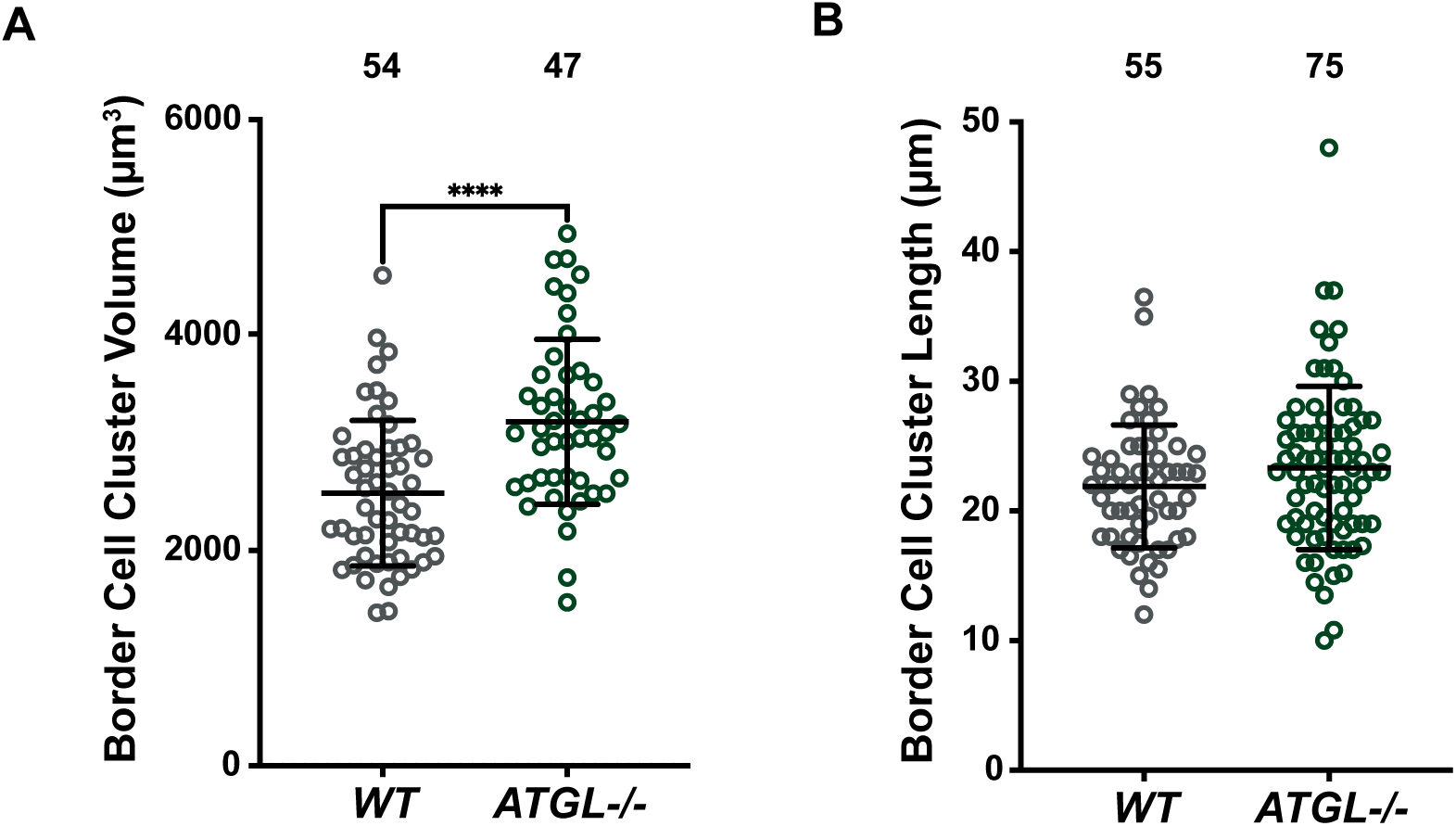
Loss of ATGL increases border cell cluster volume but does not significantly increase cluster length. **(A-B).** Quantification of the border cell cluster volume **(A)** and length **(B)** in *wild-type* (*yw*) and *ATGL* (*bmm^1^/bmm^1^*) mutants. Circle = individual border cell cluster, n = number of border cell clusters, line = mean, error bars = SD, and ns > 0.05, **** p < 0.0001, unpaired *t*-test, two-tailed. Loss of ATGL significantly increases the size of the border cell cluster **(A)** but does not increase the length of the border cell cluster **(B)**.

**Supplementary Figure 3.**
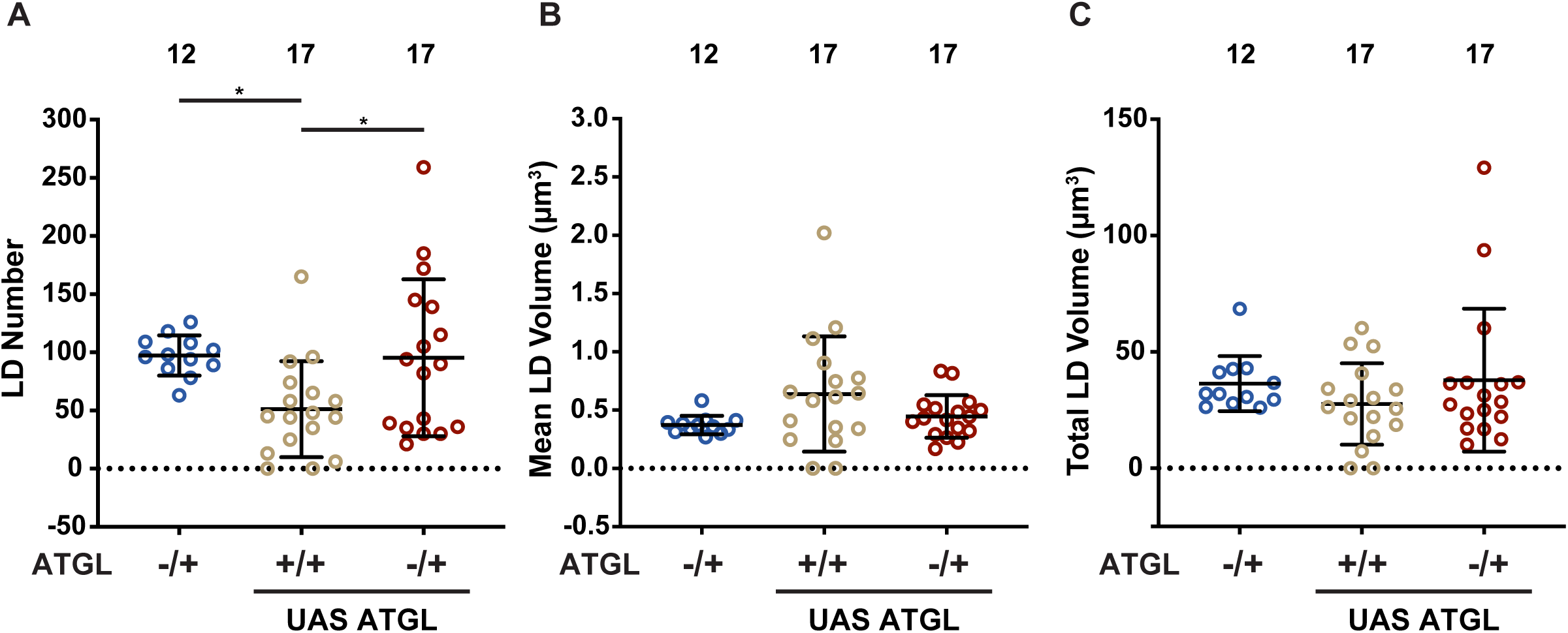
ATGL heterozygosity rescues LD numbers in UAS ATGL border cells. **(A-C).** Quantification of LD number **(A)**, mean LD volume **(B)**, and total LD volume **(C)** in the border cell clusters for the following genotypes: ATGL heterozygote (*ATGL/+*), UAS ATGL (*UAS ATGL/+*), UAS ATGL in ATGL heterozygous background (*ATGL/UAS ATGL*). Circle = individual border cell cluster, n = number of border cell clusters, line = mean, error bars = SD, and ns > 0.05, * p < 0.05, one-way analysis of variance (ANOVA) with Tukey multiple comparison test. UAS ATGL border cells have reduced LD numbers compared to ATGL heterozygotes **(A)**; however, reducing ATGL dosage (ATGL heterozygosity) restores LD numbers in UAS ATGL border cells to levels comparable to ATGL heterozygotes **(A).** Meanwhile, there are no significant differences in mean LD volume **(B)** or total LD volume **(C)** between genotypes.

**Supplementary Figure 4.**
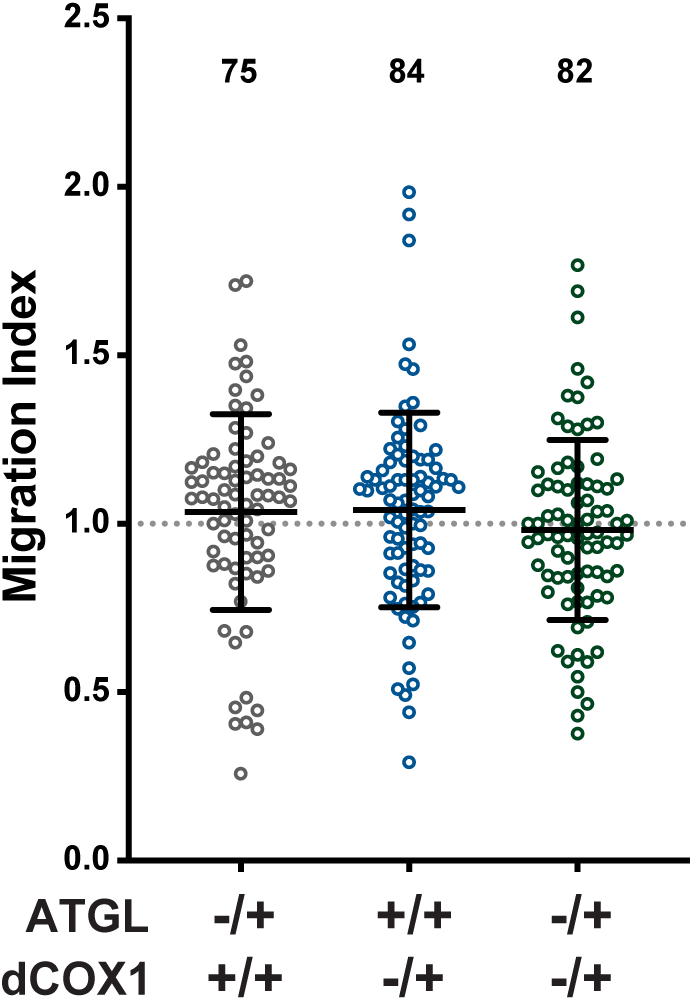
ATGL and dCOX1 do not genetically interact to promote on-time border cell migration. Graph of migration index for the following genotypes: *ATGL-/+* (*bmm^1^/+*), *dCOX1-/+* (*pxt^f01000^/+*), *ATGL-/+; dCOX1-/+* (*bmm^1^/+*; *pxt^f01000^/+*). Circle = individual border cell cluster, n = number of border cell clusters. The dotted line = on-time border cell migration, solid black lines = means, error bars = SD, and ns > 0.05, one-way analysis of variance (ANOVA) with Tukey multiple comparison test. Co-reduction of both ATGL and dCOX1 does not delay border cell migration, as the border cells from double heterozygotes of *ATGL* and *dCOX1* migrate on time.

**Supplementary Figure 5.**
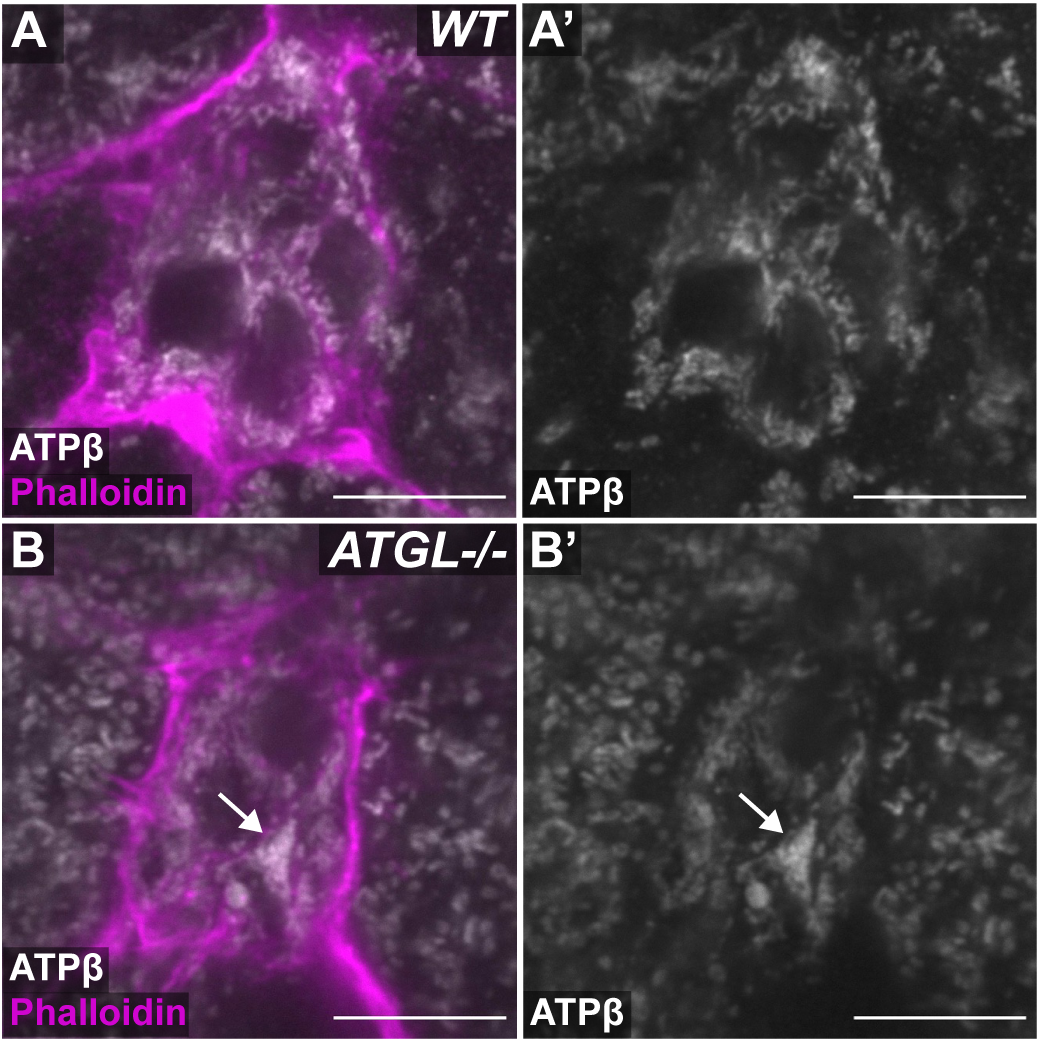
Loss of ATGL disrupts mitochondrial morphology in the border cells. **(A-B).** Maximum projections of 3 confocal slices of S9 follicles stained for mitochondria (ATPβ) in white and F-actin (Phalloidin) in magenta. White arrows indicate mitochondrial aggregates in the border cell cluster. Images brightened by 50% to increase clarity. Black boxes placed behind panel and channel labels to increase legibility. Scale bars = 10 μm. **(A-A’)** *wild-type* (*yw*). **(B-B’)** *ATGL-/-*(*bmm^1^/bmm^1^*). Loss of ATGL results in aberrant mitochondrial clustering in the border cells **(A-B’)**.

**Supplementary Figure 6.**
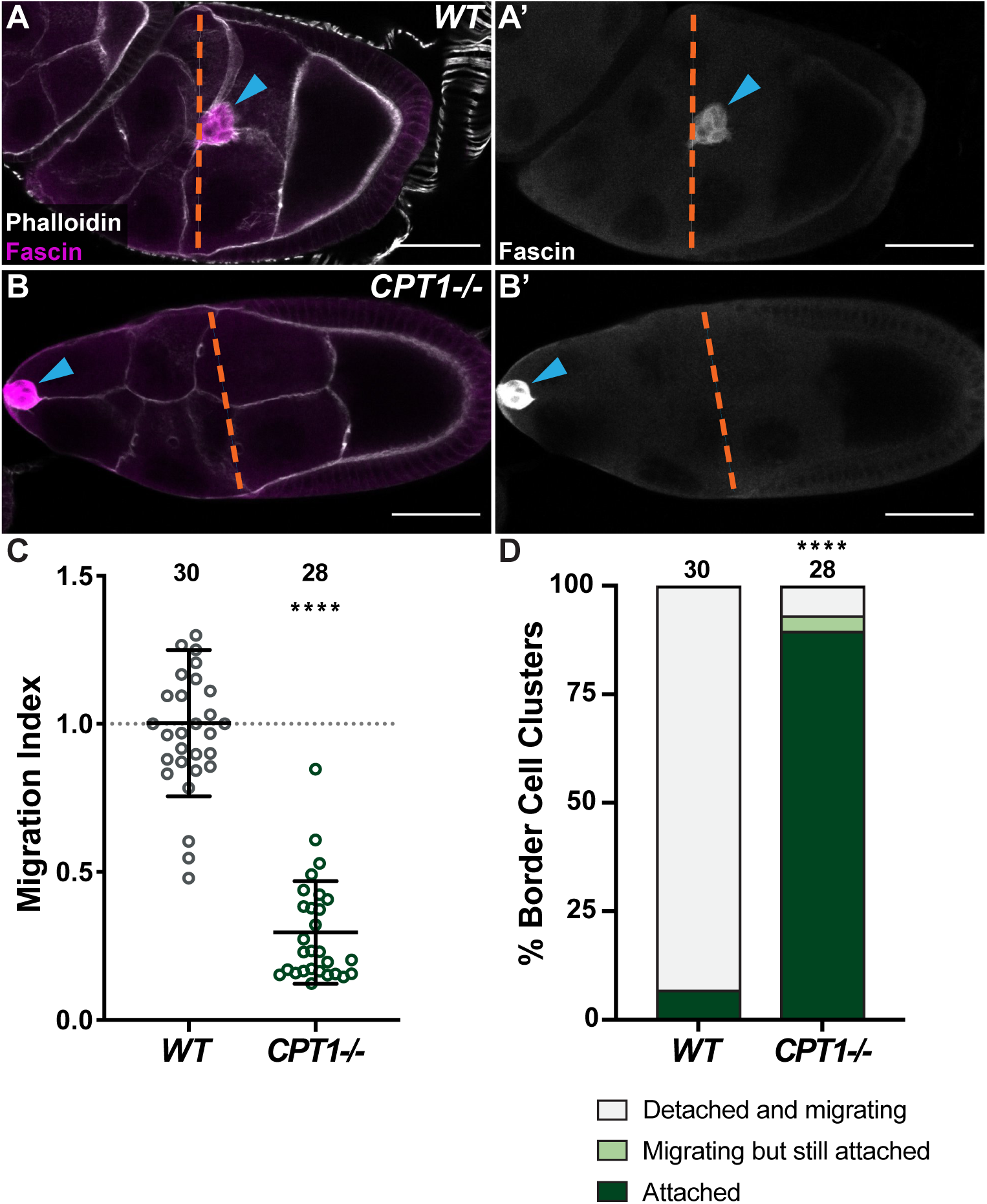
CPT1 is required for border cell detachment and on-time migration. **(A-B).** Maximum projections of 3 confocal slices of S9 follicles stained for Fascin (border cells) in magenta and F-actin (Phalloidin) in white. Blue arrowheads indicate the border cell cluster, and orange dashed lines indicate the anterior edge of the outer follicle cells. Images were not brightened. Black boxes placed behind channel labels to increase legibility. Scale bars = 50 μm. **(A, A’)** *wild-type* (*yw*). **(B, B’)** *CPT1-/-* (*whd^1^/whd^1^*). **(C).** Graph of migration index for the indicated genotypes. Circle = individual border cell cluster, n = number of border cell clusters. The dotted line = on-time border cell migration, solid black lines = means, error bars = SD, and **** p < 0.0001, unpaired *t*-test, two-tailed. **(D).** Quantification of border cell detachment defects for the indicated genotypes. Border cell clusters categorized as detached and migrating (light grey), migrating but still attached (light green), or attached (dark green). n = number of border cell clusters, **** p < 0.0001, Pearson’s Chi-squared test with Monte Carlo simulation. Loss of CPT1 delays border cell migration **(A-C)** and severely blocks detachment from the follicular epithelium **(D)**.

**Movie 1.** Loss of ATGL results in visibly larger LDs within the border cell cluster. 42 confocal slices of S9 follicles stained for LDs (BODIPY 493/503) in green and Fascin (border cells) in magenta. Movie brightened by 30% to increase clarity. Scale bars = 10 μm. Left: *wild-type* (*yw*). Right: *ATGL-/-*(*bmm^1^/bmm^1^*). Loss of ATGL results in larger LDs within the border cell cluster relative to wild-type.

**Supplementary Table S1: Reagents used in the study.**

**Supplementary Table S2: List of genotypes by figure.**

**Supplementary Table S3: Raw data from the study.**

